# Community composition shapes microbial-specific phenotypes in a cystic fibrosis polymicrobial model system

**DOI:** 10.1101/2022.06.23.497319

**Authors:** Fabrice Jean-Pierre, Thomas H. Hampton, Daniel Schultz, Deborah A. Hogan, Marie-Christine Groleau, Eric Déziel, George A. O’Toole

## Abstract

Interspecies interactions can drive the emergence of unexpected microbial phenotypes that are not observed when studying monocultures. The cystic fibrosis (CF) lung consists of a complex environment where particular microbes, living as polymicrobial biofilm-like communities, are associated with negative clinical outcomes for persons with CF (pwCF). However, the current lack of *in vitro* models integrating the microbial diversity observed in the CF airway hampers our understanding of why polymicrobial communities are recalcitrant to therapy in this disease. Here, integrating computational approaches informed by clinical data, we built a mixed community of clinical relevance to the CF lung composed of *Pseudomonas aeruginosa, Staphylococcus aureus, Streptococcus sanguinis* and *Prevotella melaninogenica*. We developed and validated this model biofilm community with multiple isolates of these four genera. When challenged with tobramycin, a front-line antimicrobial used to treat pwCF, the microorganisms in the polymicrobial community show altered sensitivity to this antibiotic compared to monospecies biofilms. We observed that wild-type *P. aeruginosa* is sensitized to tobramycin in a mixed community versus monoculture, and this observation holds across a range of community relative abundances. We also report that LasR loss-of-function, a variant frequently detected in the CF airway, induces tolerance of *P. aeruginosa* to tobramycin specifically in the mixed community. The molecular basis of this community-specific recalcitrance to tobramycin for the LasR mutant variant is the increased production of redox-active phenazines. Our data support the importance of studying clinically-relevant model polymicrobial biofilms to understand community-specific traits relevant to infections.

**SIGNIFICANCE STATEMENT:** The CF lung is colonized by biofilm-like microbial communities that exhibit both resistance and tolerance (collectively called “recalcitrance”) to antimicrobials used in the clinic. Here, we leveraged clinical data from pwCF to inform our understanding of communities exhibiting recalcitrance. We developed and validated an *in vitro* model that revealed novel, community-specific phenotypes relevant to the clinic. We used this model to explore the underlying mechanism associated with a community-specific emergent behavior. We posit that *in vitro* models of polymicrobial communities may help in developing new antimicrobial strategies to improve patient outcomes, and that the approach used here can be applied to other polymicrobial models.

## INTRODUCTION

The disconnect between *in vitro* antimicrobial susceptibility profiles and clinical response poses a significant threat to the eradication of mixed microbial communities observed in human infections (1). That is, while minimal inhibitory concentration (MIC) assays are widely used to guide clinical intervention by determining the antimicrobial susceptibility profiles of pathogenic species, such approaches often fail to resolve chronic human-based, polymicrobial infections (2, 3). Studies published several decades ago documented that microbe-microbe interactions can drive shifts in antimicrobial susceptibility profiles (4, 5). Thus, antibiotics effective at killing a single microorganism *in vitro* may be ineffective against the same microbe when grown in the context of a polymicrobial infection. More recent studies provide some insight as to the mechanisms driving resistance and tolerance (also referred to as recalcitrance or resilience), towards antimicrobials in the context of mixed-species infections (6-11). These mechanisms include (but are not limited to) the production of metabolites altering microbial physiology, horizontal gene transfer of genetic material, and passive protection by the production of shared “public goods”, such as β-lactamases (reviewed in (2, 12)).

Over the last decade, both culture-based and culture-independent studies have indicated that chronic cystic fibrosis (CF) lung infections are characterized by the presence of numerous microbial taxa (13-16). The clinical-relevance of such mixed communities is highlighted by the observations that co-occurrence of *Pseudomonas aeruginosa* and *Staphylococcus aureus* alters the antimicrobial susceptibility of the latter organism and worsens CF lung disease (9-11, 17). Furthermore, antibiotics effective against pathogens such as *P. aeruginosa* in classic *in vitro* MIC assays show limited clinical efficacy, as indicated by the inability of tobramycin to clear this microorganism from the CF airway (18-20). Indeed, a recent report recommends abandoning MIC testing for microorganisms isolated from CF airway infections as their predictive value for a positive treatment outcome is not supported (3).

The pathogenesis of mixed-species infections in the CF airway is still poorly understood but it is now appreciated that distinct community types can impact clinical outcomes (21, 22). However, one of the current missing links allowing for the translation of microbiome-informed studies back to the clinic is the establishment of *in vitro* mixed community models that can be used to understand community function(s) of CF pathogens (23). That is, although some CF-relevant model communities have been proposed (9-11, 24-26), they (i) do not entirely reflect the polymicrobial nature of the CF airway and/or (ii) do not utilize *in vitro* conditions recapitulating the nutritional milieu and/or biofilm-like growth observed in the CF lung. Thus, there is a pressing need for the development of clinically-informed, *in vitro* polymicrobial communities to probe the molecular mechanism(s) governing microbial interactions, particularly in regard to the responsiveness to antimicrobial agents.

The aims of this study were four-fold: (i) to leverage large, existing microbiome data sets and their associated clinical metadata to inform and develop a clinically-relevant and tractable *in vitro* model of the polymicrobial communities found in the CF airway, (ii) to identify community-specific phenotypes, (iii) to test this *in vitro* model system against the most common front-line CF antimicrobial, tobramycin (27), and (iv) to provide a mechanistic understanding whereby community composition can impact tobramycin responsiveness.

## RESULTS

### Using computational approaches to identify community types found in the CF airway

We sought to exploit available 16S rRNA gene amplicon library sequencing data and associated clinical metadata to identify a representative set of microbial communities found in persons with CF (pwCF) that we could model *in vitro*. A recent study from our team using k-means to cluster 16S rRNA gene amplicon sequencing relative abundance values and another machine learning approach (e.g., the gap statistic) identified five clusters as the most parsimonious number (21). This same study revealed, following the analysis of a large cross-sectional 16S rRNA gene sequence data sets and associated metadata, that the 5 community types explained 24% of variability in lung function, twice as high as any other factor or group of factors previously identified (21). Thus, two different approaches, using clinical outcomes data, identified five similar community types.

Of these five communities, two were dominated by a single microbial taxa: *P. aeruginosa* (designated the “Pa.D” community) and *Streptococcus* (Strep.D); these communities included 73/167 (∼23%) of the analyzed samples (21). Several *P. aeruginosa*-focused CF studies make it clear that specifically targeting this pathogen does not translate to positive clinical outcomes in pwCF (18, 19, 28, 29) and there has already been a substantial focus on the study of *P. aeruginosa* in monoculture in the context of CF (30, 31). Thus, we deemed that *in vitro* modeling of a *P. aeruginosa*-dominated community would not allow us to probe unknown/novel factors driving chronic CF lung disease. Furthermore, while the presence of *Streptococcus* spp. may influence CF airway health, its clinical relevance in CF is still a matter of active research and this genus is associated with both worsened and improved clinical outcomes (32). We identified a third cluster designated Oth.D, for “other”; this cluster was comprised of samples from pwCF whose lung microbiota were dominated by less-common, individual genera including *Stenotrophomonas, Burkholderia* and *Achromobacter*. Based on these factors, we decided to focus on two community types identified in the study cited above (21), which are composed of various microbial taxa (Pa.M1/Pa.M2; which we refer to as “mixed”). The Pa.M1 and Pa.M2 mixed communities represented ∼50% of the analyzed samples in this study (21).

Using the known relative abundance of *P. aeruginosa, S. aureus, Streptococcus sanguinis* and *Prevotella melaninogenica* in each of these two mixed (Pa.M1/Pa.M2) communities, we performed modeling of metabolic fluxes and compared these findings to metabolic fluxes with *Pseudomonas*-dominated (Pa.D) and *Streptococcus*-dominated (Strep.D) communities. We noted multiple similarities between the predicted metabolic fluxes of Pa.M1 and Pa.M2 that distinguished these mixed communities from the *Pseudomonas*- and *Streptococcus*-dominated conditions (**Fig. S1A**). Thus, we considered these two mixed communities as functionally similar and decided to focus on the development of a single “mixed” community model.

Next, we sought to identify a limited number of community members to model as a tractable *in vitro* mixed community. We noted that ten microbial taxa achieved relatively high abundance and prevalence across the five identified community types (21) (**Fig. 1A**). From among these taxa we decided to focus on *P. aeruginosa, S. aureus, Streptococcus* spp. and *Prevotella* spp. as members of our model mixed community for the following reasons: (i) *P. aeruginosa, S. aureus, Streptococcus* spp. and *Prevotella* spp. are important in shaping health outcomes in pwCF based on studies using culture-dependent and -independent approaches (30, 33-39). (ii) Additional studies provide evidence of the potential for two or more of these microorganisms found together in the CF airway to lead to worsened clinical outcomes (17, 40, 41). (iii) Imaging studies of sputum have shown evidence for a sub-set of these microbes being in close physical proximity (42, 43). (iv) Assessing the fraction of 16S rRNA reads across the data sets analyzed by Hampton, O’Toole and colleagues (21), *Pseudomonas, Staphylococcus, Streptococcus* and *Prevotella* were among the top 10 microbial taxa detected in the CF airway with relative abundances of 35%, 3%, 20% and 13% and prevalence rates of 90%, 50%, 80% and 70%, respectively (21) (**Fig. 1A**). (v) In 103 of the 167 (∼62%) patient samples analyzed by Hampton, O’Toole and colleagues, ≥70% of 16S rRNA gene reads were assigned to *Pseudomonas, Staphylococcus, Streptococcus* and *Prevotella* (**Fig. 1B**); another 30 samples contained at least 50% of the reads assigned to these four genera. Including the next most abundant and prevalent genera such as *Burkholderia* or *Achromobacter* as a 5^th^ member of the mixed community would only marginally increase (by ∼6%) the number of patient samples covered (**Fig. S1B**,**C**). Furthermore, the average abundance of *Burkholderia* was skewed by a relatively small number of patients in our cohort that were dominated by this pathogen (21). (vi) Finally, work published by our group using metabolic modeling indicates that *P. aeruginosa, S. aureus, S. sanguinis* and *P. melaninogenica* are top contributors of cross-fed metabolites in communities detected in the CF lung (44). That is, their abundance in the CF airway could be explained by predicted metabolic cross-feeding among these four microorganisms. Thus, we settled on a mixed community comprised of *P. aeruginosa, S. aureus, Streptococcus* spp. and *Prevotella* spp., which represents a common “pulmotype” detected in ∼50% of airway infections for pwCF, as the basis for the development of an *in vitro* model system.

**Figure 1.**
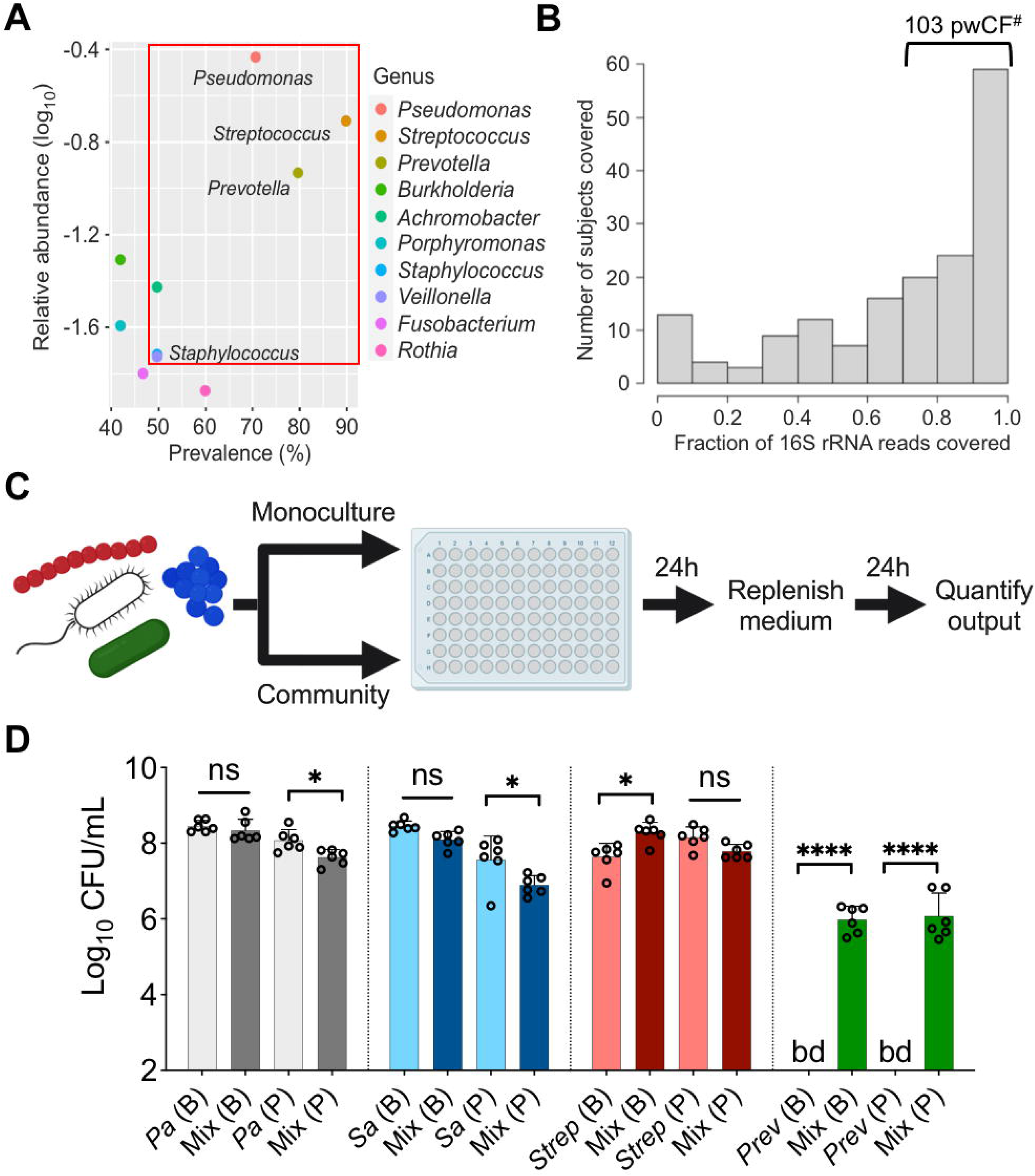
Leveraging clinical microbiome data sets with computational analyses to identify communities and community members to model *in vitro*. (**A**) Relative 16S rRNA gene abundance and prevalence of the top 10 CF lung pathogens in the 167 pwCF data set used as the basis for developing the *in vitro* mixed community, as reported (18). (**B**) Number of unique samples for which ≥70% of 16S rRNA reads are associated to the combined presence of *Pseudomonas, Staphylococcus, Streptococcus, Prevotella*. ^*#*^Indicates the number of samples (103) that meet this criterion from the total sample size of 167 pwCF. (**C**) Experimental design for the cultivation of monoculture versus mixed community cultures in a plastic 96-well plate. (**D**) Colony forming units (CFU) counts of each microbial member grown as a monoculture and in a mixed community (Mix) for biofilm (B) and planktonic (P) fractions. CFU were performed by plating on medium selective for the growth of each microorganism. Each column represents the average from at least three biological replicates, each with at least three technical replicates. Statistical analysis was performed using ordinary one-way analysis of variance (ANOVA) and Tukey’s multiple comparisons posttest with *, *P* < 0.05; ****, *P* < 0.0001, ns = non-significant. *Pa* = *Pseudomonas aeruginosa, Sa* = *Staphylococcus aureus, Strep = Streptococcus* spp., *Prev* = *Prevotella* spp., bd = below detection.

### Implementing a mixed *in vitro* model system to probe community function

Based on the data outlined in the previous section, we focused on four bacteria: *P. aeruginosa, S. aureus, Streptococcus* spp. and *Prevotella* spp. We cultivated laboratory strains of *P. aeruginosa* PA14, *S. aureus* Newman, *S. sanguinis* SK36 and *P. melaninogenica* ATCC25845 as monocultures and mixed communities in anoxic conditions (**Fig. 1C**), which reflects the CF airway environment (45), and using artificial sputum medium (ASM), which mimics the nutritional conditions of the CF lung (46, 47). We could quantify the respective viable counts from both biofilm and planktonic populations for each of these four microorganisms by plating on selective media, as shown in **Figure 1D** and described below.

While the *P. aeruginosa* biofilm population did not show statistically significant differences in endpoint CFU counts at 24 h when cultivated as either a monoculture or in a mixed community, a modest (∼0.5 log) but statistically significant decrease was detected for *P. aeruginosa* grown planktonically in the mixed community (**Fig. 1D**, grey bars). While not negatively impacted in a mixed community versus monoculture in a biofilm, *S. aureus* viable counts were reduced in a planktonic community versus when cultivated as monoculture (**Fig. 1D**, blue bars). A statistically significant increase in biofilm CFU counts was observed for *S. sanguinis* in the mixed community (**Fig. 1D**, red bars). However no differences were detected for planktonic cells of this microorganism in the presence of microbial partners (**Fig. 1D**, red bars). Interestingly, we could not detect *P. melaninogenica* in monoculture, a finding predicted in our previous metabolic modeling study (44), but ∼6 log_10_ CFU/mL of this microorganism could be detected when cultivated in the presence of other microbial partners in both biofilm and planktonic fractions (**Fig. 1D**, green bars).

The *in vitro* mixed biofilm community under the model growth conditions resulted in a composition within the range observed for the M1/M2 mixed communities previously reported (21) (**Fig. S2A**). Moreover, by maintaining *P. aeruginosa* at the same starting concentration and shifting the inoculum of *S. aureus, S. sanguinis* and *P. melaninogenica*, different mixed community compositions reflecting the microbial populations in the CF lung were observed (**Fig. S2B**).

While the data shown in **Figure 1D** pertains to commonly used laboratory strains, similar observations were made for multiple other strains and/or CF clinical isolates for these four microbial genera (**Fig. S3**). Finally, we assessed whether this model system could maintain these microbial populations over time by replacing the medium every 24 hours. We stably detected each member of the mixed community for up to 2 weeks (**Fig. S4**). Taken together, our data shows that we can model an *in vitro* mixed community reflective of the polymicrobial infection in the CF airway found in the airway of ∼50% of pwCF, and that the microorganisms in this model display community-specific growth phenotypes.

### Polymicrobial context shifts tobramycin sensitivity of CF pathogens

Using the newly developed model described above, we sought to test the hypothesis that the susceptibility of CF pathogens to tobramycin would shift when treated in the context of a mixed community. Tobramycin was selected as it is one of the most heavily prescribed antimicrobials in the CF clinic (27); we used clinically relevant concentrations in all studies (48).

Initially focusing on biofilm communities, which are more recalcitrant to antimicrobial therapy (49), we observed that tobramycin treatment of wild-type (WT) *P. aeruginosa* PA14 grown in the mixed community resulted in an unexpected reduction in the number of viable cells compared to monoculture (**Fig. 2A**). Furthermore, no detectable counts of *P. aeruginosa* in the planktonic phase of the mixed community were observed (**Fig. 2A**). This sensitization of *P. aeruginosa* was reproducible in the presence of a wide range of microbial partners grown in the mixed community (**Fig. S5**). Notably, inoculating the community with a 1000X fold less CFU of *S. aureus, S. sanguinis* and *P. melaninogenica* abrogated the community-dependent enhanced tobramycin killing against *P. aeruginosa* (**Fig. S5**). These data demonstrate increased killing of *P. aeruginosa* by tobramycin in a polymicrobial environment can occur over a range of abundances of other microbial partners, but there is a lower limit to the presence of other partners to observe community-specific phenotypes.

**Figure 2.**
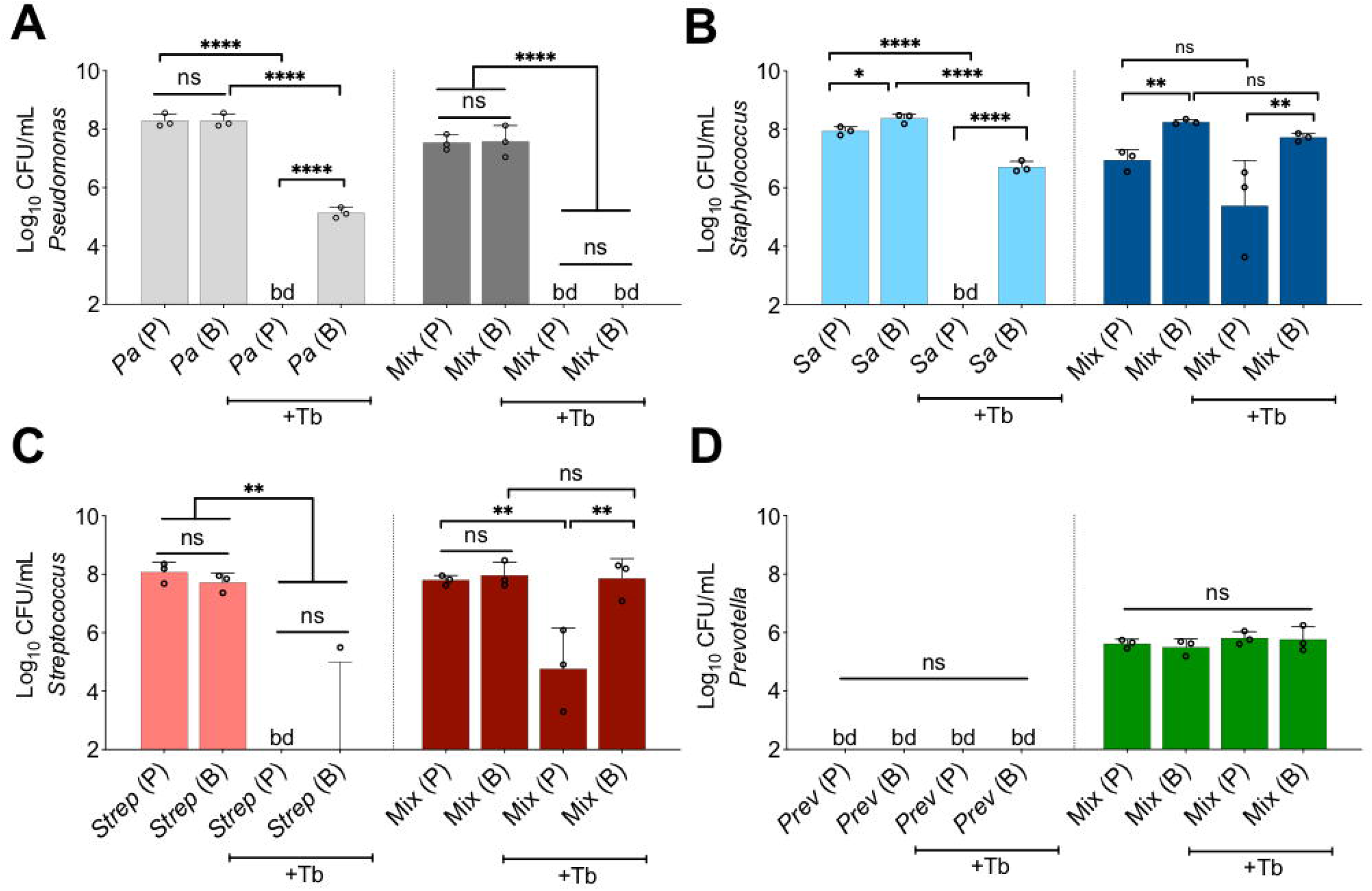
Polymicrobial context shifts tobramycin sensitivity of CF pathogens. Colony forming units of planktonic (P) and biofilm (B) populations of (**A**) *P. aeruginosa* (*Pa*), (**B**) *S. aureus* (*Sa*), (**C**) *S. sanguinis* (*Strep*) and (**D**) *P. melaninogenica* (*Prev*) grown as monocultures or mixed communities (Mix) and challenged or not with 100 µg/mL of tobramycin (+Tb). Each column represents the average from at least three biological replicates, each with at least three technical replicates. Statistical analysis was performed using ordinary one-way analysis of variance (ANOVA) and Tukey’s multiple comparisons posttest with *, *P* < 0.05; **, *P* < 0.01; ****, *P* < 0.0001, ns = non-significant. bd = below detection.

We expanded our analysis beyond the PA14 laboratory strain of *P. aeruginosa*. Of the six *P. aeruginosa* CF clinical isolates tested (including mucoid and non-mucoid strains), two strains displayed a similar phenotype to strain PA14 (**Fig. S6**: strains SMC1596, AMT0101-1-2, labeled in red), one strain showed equal sensitivity in monoculture and the polymicrobial community (**Fig. S6**: strain SMC1587), while one clinical isolate displayed increased tolerance in a polymicrobial environment (**Fig S6**: strain SMC1595, labeled in blue). A similar enhancement of tobramycin-mediated killing was observed when *P. aeruginosa* PA14 was co-cultivated with multiple *Streptococcus* and *Prevotella* species (**Fig. S7**). Interestingly, co-cultivating *P. aeruginosa* PA14 with *S. aureus* strain USA300 did not impact tobramycin sensitivity of *P. aeruginosa* in the mixed community while strains Newman and JE2 did alter sensitivity (**Fig. S7C**), indicating that *S. aureus* might contribute to the community-specific shift in *P. aeruginosa* sensitivity to tobramycin.

We noted several other changes in tobramycin sensitivity in the context of the mixed community compared to monoculture. Most *S. aureus* strains and *Streptococcus* spp. showed decreased tobramycin susceptibility when cultivated in the mixed community in both planktonic and biofilm populations (**Figs. 2B,C, S8, S9**). *S. aureus* strain USA300 showed high level tolerance to tobramycin in monoculture, but was not further protected from tobramycin in the mixed community (**Fig. S8A**). We could not determine shifts in *Prevotella* spp. sensitivity to tobramycin in monoculture as this microorganism cannot be cultivated under these conditions in the absence of the other community members (**Figs. 2D** and **S10**). Community-specific protection of bacterial members was unlikely driven by microbial-based inactivation of the drug as the minimal bactericidal concentration (MBC) value remained similar to ASM control when *P. aeruginosa* was treated with tobramycin that was pre-incubated with microbial supernatants for 24 h (**Table S1**). Taken together, we demonstrated that growth in the mixed community shifts the antimicrobial sensitivity of multiple bacteria compared to growth in monoculture.

### LasR loss of function increases tobramycin tolerance of *P. aeruginosa* in the mixed community

The unexpected increase in sensitivity of *P. aeruginosa* when challenged with tobramycin in the mixed community (**Fig. 2A**) prompted us to reconcile our *in vitro* observations with what is observed in the clinic. That is, culture-independent microbiome studies do not indicate any appreciable changes in the abundance of *P. aeruginosa* in the airways of pwCF post-treatment with tobramycin (18-20).

*P. aeruginosa* undergoes genetic adaptation in the airway of pwCF (50) and loss-of-function mutations in the gene coding for the key intercellular communication regulator, LasR (51), are frequently observed in the airway of pwCF and other environments (52-56). Strains lacking LasR function have also been associated with worsened lung function (54), *in vitro* growth advantage in low oxygen conditions (57), increased production of virulence factors through intraspecies interactions (58) and greater tolerance to front-line CF drugs (59). We thus hypothesized that loss of LasR function might result in a different response towards tobramycin treatment in the community.

To test this hypothesis, we replaced WT *P. aeruginosa* PA14 in the mixed population with an isogenic Δ*lasR* mutant. The absence of LasR function resulted in increased tolerance to tobramycin in the mixed community (**Fig. 3A**). This phenotype was complemented by restoring a WT *lasR* allele at the native locus (**Fig. 3A**). Importantly, the Δ*lasR* mutant showed a similar degree of sensitivity to tobramycin as the WT when these strains were grown in monoculture (**Fig. 3A**). The recalcitrance of the Δ*lasR* mutant was observed to be biofilm-specific as planktonic cells grown in a mixed community displayed equal sensitivity to the drug when compared to monoculture (**Fig. 3B**). Endpoint CFU counts of other microbial members in the mixed community were not impacted by the inactivation of the *lasR* gene with or without tobramycin treatment (**Fig. S11**). Inactivation of tobramycin by the Δ*lasR* mutant was unlikely as the *P. aeruginosa* minimal bactericidal concentration (MBC) remained similar to ASM control when WT *P. aeruginosa* was treated with tobramycin pre-incubated in the supernatant of the Δ*lasR* mutant (**Table S1**).

**Figure 3.**
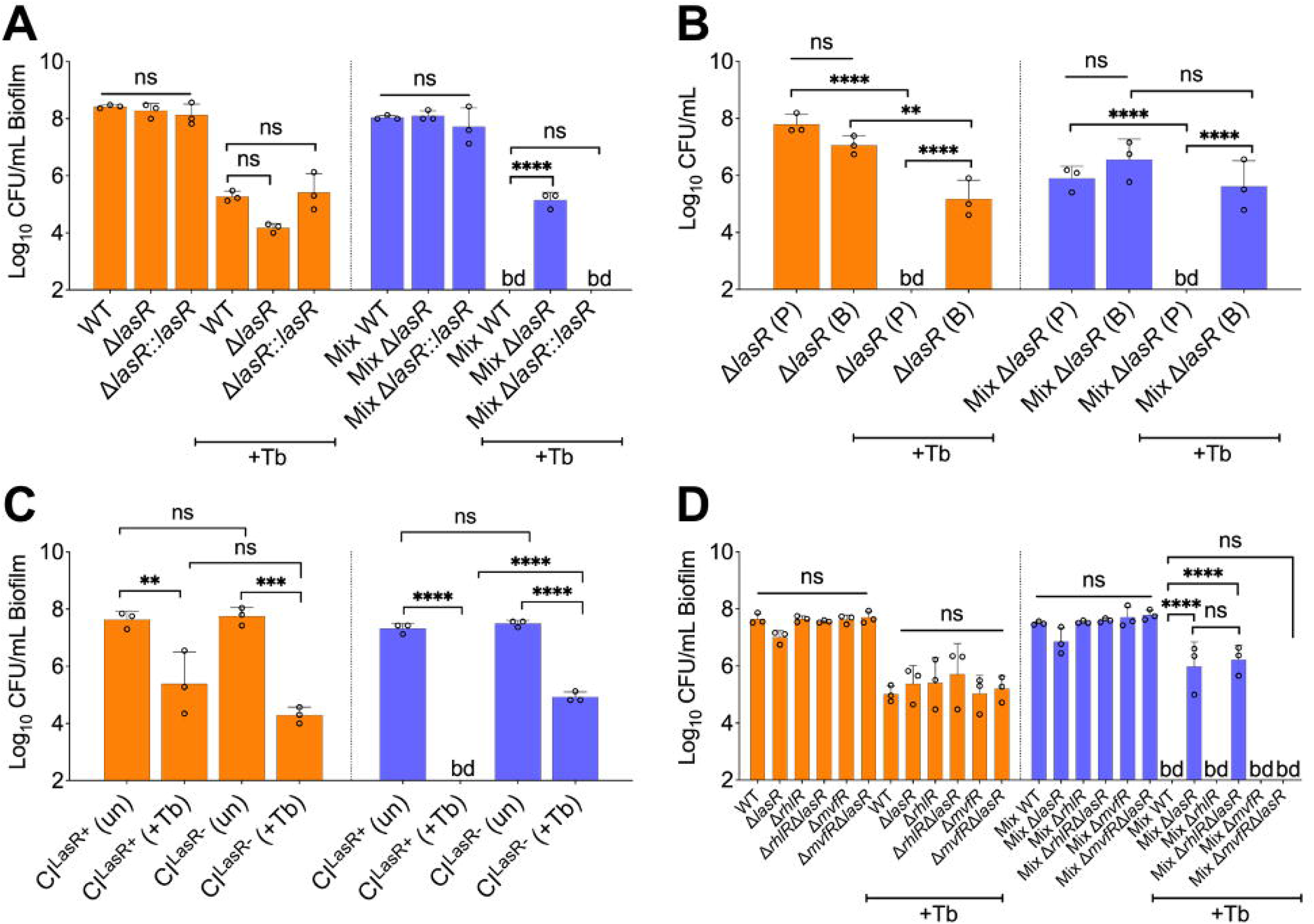
LasR loss-of-function drives biofilm-specific tobramycin tolerance in a mixed community. Colony forming units of (**A**) *P. aeruginosa* strain PA14 (WT), isogenic Δ*lasR* mutant and the complemented strain (Δ*lasR*::*lasR*), (**B**) planktonic and biofilm Δ*lasR* mutant cells, (**C**) LasR-defective (NC-AMT0101-1-1; LasR-) and LasR+ (NC-AMT0101-1-2) clinical isolates (CI) and (**D**) *P. aeruginosa* quorum sensing regulator mutants grown as monocultures and mixed communities (Mix) and challenged or not with 100 µg/mL of tobramycin (+Tb). Each column represents the average from at least three biological replicates, each with at least three technical replicates. Statistical analysis was done using ordinary one-way analysis of variance (ANOVA) and Tukey’s multiple comparisons posttest with **, *P* < 0.01; ***, *P* < 0.001; ****, *P* < 0.0001. bd = below detection, un = untreated.

To determine whether the altered tolerance of LasR loss-of-function is not specific to the *P. aeruginosa* PA14 strain, we tested a chronic CF clinical isolate, NC-AMT0101-1-1, defective for LasR activity (LasR^-^) (50). The strain displayed higher tolerance to tobramycin in the mixed community than its parent isolate (NC-AMT0101-1-2; LasR^+^) (**Fig. 3C**). Both NC-AMT0101-1-1 and NC-AMT0101-1-2 isolates were equality sensitive to tobramycin in monoculture suggesting a community-specific phenotype for LasR loss-of-function in the model (**Fig. 3C**). These results further prompted us to test the hypothesis that the clinical isolate SMC1595 lacked LasR function as this strain is more tolerant to tobramycin treatment in a mixed community (**Fig. S6**). Indeed, both low protease activity and reduced 3-oxo-C_12_ homoserine lactone production, features associated with LasR loss-of-function variants (52, 53, 55), were observed in that strain when compared to controls (**Fig. S12**).

The LasR quorum sensing (QS) regulator influences the function of many target systems in *P. aeruginosa* including the QS regulators RhlR and MvfR (also known as PqsR) (51). We tested the impact of the inactivation of RhlR or MvfR in a Δ*lasR* mutant background to understand how LasR influences community tolerance. We observed that recalcitrance of the Δ*lasR* mutant in the mixed community is dependent on the MvfR-PQS pathway, as the absence of a functional MvfR regulator resensitized the Δ*lasR* mutant to tobramycin (**Fig. 3D**). On the other hand, the inactivation of RhlR QS regulator, did not impact the sensitivity of the Δ*lasR* mutant (**Fig. 3D**). None of the tested mutants, including Δ*lasR*, Δ*rhlR*, Δ*mvfR*, Δ*mvfR*Δ*lasR*, Δ*rhlR*Δ*lasR*, resulted in a change in the endpoint CFUs of the other members in the mixed community in the presence or absence of tobramycin (**Fig. S13**). Our data indicate that the tolerance of the Δ*lasR* mutant is biofilm-specific and implicates the MvfR-PQS QS system in this increased tolerance observed for this mutant in the mixed community.

### Phenazines induce *P. aeruginosa* tolerance in the mixed community

The LasR regulon includes genes associated with the production of multiple virulence factors, including the redox-active molecules phenazines (60). The LasR-regulated MvfR-PQS system also impacts phenazines production (61, 62). Phenazines have previously been shown to drive tolerance of *P. aeruginosa* to various drug classes, including aminoglycosides (63-65). Based on our observations (**Fig. 3**) and given that loss of LasR function often results in overproduction of phenazines in various contexts (53, 58, 66-68), we hypothesized that these molecules might be overproduced in the mixed community thereby conferring tolerance to the Δ*lasR* mutant. To test this hypothesis, we quantified, phenazine-1-carboxylic acid (PCA) produced by wild-type and associated mutants of *P. aeruginosa*. Indeed, we measured significantly higher PCA levels in the mixed community containing the Δ*lasR* mutant when compared to WT *P. aeruginosa* (**Fig. 4A**).

**Figure 4.**
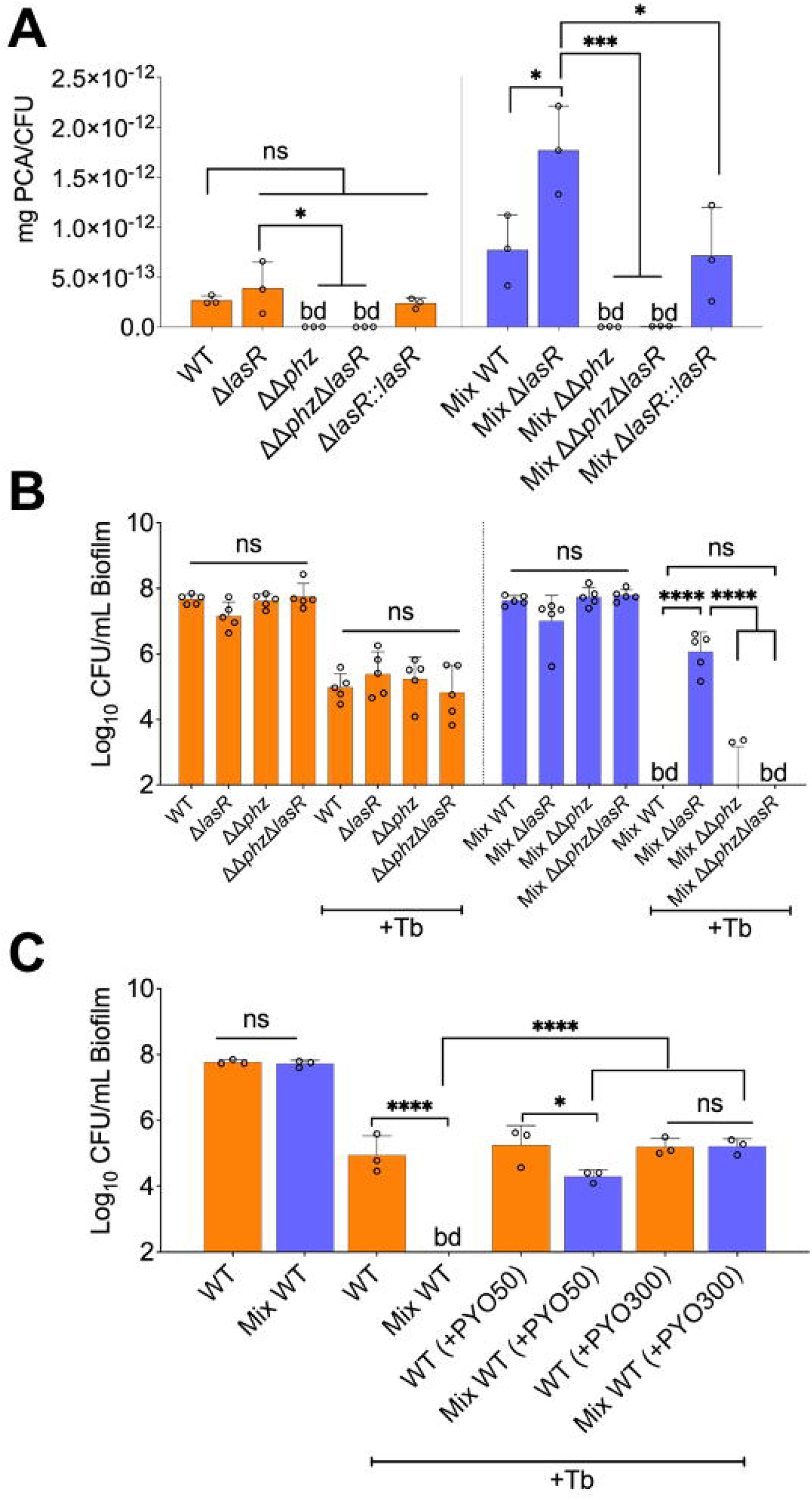
Phenazines drive tolerance of *P. aeruginosa* in mixed communities. (**A**) HPLC-MS/MS quantification of phenazines in monoculture and mixed communities (Mix) containing the indicated *P. aeruginosa* WT and mutant strains. (**B**) Colony forming units (CFU) counts of *P. aeruginosa* (WT) and indicated mutants grown as monoculture or mixed communities (Mix). (**C**) Exogenous addition of phenazine to monoculture and mixed communities with wild-type *P. aeruginosa* treated with 100 µg/mL tobramycin (+Tb). Two physiologically-relevant phenazine concentrations were tested; 50 µM (+PYO50) and 300 µM (+PYO300). Data for *S. aureus, S. sanguinis* and *P. melaninogenica* counts are shown in **Figure S15**. Each column represents the average from at least three biological replicates, each with at least three technical replicates. Statistical analysis was performed using ordinary one-way analysis of variance (ANOVA) and Tukey’s multiple comparisons posttest with * = *P* < 0.05; *** = *P* < 0.001; **** = *P* < 0.0001, ns = non-significant, bd = below detection.

Also consistent with our hypothesis, blocking the production of phenazines in the Δ*lasR* background (ΔΔ*phz*Δ*lasR*, which lacks both *phz* operons) resensitized the Δ*lasR* mutant strain to tobramycin in the mixed community (**Fig. 4B**). None of the tested mutants impacted the abundance of other members in the mixed community in the presence or absence of tobramycin (**Fig. S14A**). Based on the findings using mutants, we sought to test the hypothesis that exogenous addition of phenazine induces tolerance of *P. aeruginosa* in the mixed community. Treatment of pre-formed biofilms with the phenazine pyocyanin triggered tolerance of WT *P. aeruginosa* in the mixed community in a dose-dependent manner (**Fig. 4C**). The tested phenazine concentration did not impact endpoint CFUs of *P. aeruginosa* in all conditions tested (**Fig. S14B**).

## DISCUSSION

By leveraging culture-independent studies in combination with clinical metadata, we built a stable CF-relevant polymicrobial model system composed of *P. aeruginosa, S. aureus, S. sanguinis* and *P. melaninogenica* (**Fig. 1**). We used experimental conditions reflecting the nutritional (e.g., artificial sputum medium) and environmental nature (e.g., anoxia) of the CF airway (42, 45, 46). The community fell within the range of clinically observed communities in terms of species relative abundance (**Fig. S2A**), and furthermore, we observed several community-specific phenotypes (**Fig. 1D, Fig. S3**). In agreement with previous reports (69-72), *S. aureus* and *Streptococcus* spp. showed decreased and increased growth, respectively, in the mixed community versus monoculture. While other studies have demonstrated that *P. aeruginosa*-secreted exoproducts can eradicate *S. aureus* in co-culture (69, 73), we could maintain *S. aureus* viability in the mixed community for up to 14 days under the CF-like conditions used here (**Fig. S4**). Thus, the observations in our polymicrobial community mirror the capacity of these two pathogens to co-exist in the CF airway (74).

A major hurdle in the eradication of microbial communities detected in the CF airway is their resilience to front-line CF antimicrobials (2, 12). That is, while pathogens tested in monoculture demonstrate sensitivity to several antimicrobial classes (2, 12), these findings do not translate to improved outcomes in the clinic (18, 19, 29). For example, a recent report shows that current testing methods fail to predict outcomes in pwCF (3). By modeling a polymicrobial community of six frequently encountered pathogens in the CF lung, Vandeplassche and colleagues compared the impact of several CF drugs on microbes grown as monocultures versus in a mixed community (26). Surprisingly, no shifts in sensitivities of any of the microorganisms were observed. These results could perhaps be explained by the differences in experimental conditions as Vandeplassche and colleagues used a rich medium and incubations were performed under normoxic conditions (26). However, we acknowledge that this study represents an important first step in tackling the impact of microbial interactions on antimicrobial susceptibility of CF pathogens.

Using more “CF-like” experimental conditions informed by existing clinical data sets, we sought to test the hypothesis that growth in a mixed community would shift the sensitivity of CF microorganisms versus growth as monocultures. We focused on the front-line CF antimicrobial tobramycin as (i) this drug is the most heavily prescribed therapeutic to treat chronic CF lung infections (27) but (ii) often does not eradicate pathogens (especially *P. aeruginosa*) from the airways of pwCF (18, 19). Using our system, we made several observations. First, almost all tested *S. aureus* and *Streptococcus* spp. strains were protected from tobramycin-mediated killing in the mixed community versus monoculture (**Fig. 2B,C, S8, S9**). In a recently published study, Murray and colleagues observed that *S. aureus* can be protected from tobramycin eradication in the presence of *P. aeruginosa* (75), but this protection was abolished in the presence of a PQS operon inhibitor (75). Inactivation of the MvfR/PqsR regulator, which controls the expression of the PQS operon (61), did not sensitize *S. aureus* to tobramycin under our experimental conditions (**Fig. S11**) indicating that other mechanisms might be driving this phenotype in our system. Furthermore, the observed increased tobramycin resilience of *Streptococcus* spp. in a polymicrobial environment (**Figs. 2D, S9**) is in agreement with a previous report (76). Future studies will be necessary to identify the community-specific recalcitrance mechanisms employed by *S. aureus* and *Streptococcus* spp.

The decrease in the viability of *P. aeruginosa* in a tobramycin-treated community (**Fig. 2A, S5, S6 and S7**) was unexpected. That is, recent culture-independent studies investigating the impact of tobramycin in pwCF do not show any appreciable decrease in *P. aeruginosa* after extended exposure to this antibiotic (18, 19). These results prompted us to interrogate the molecular mechanisms driving this increased sensitivity of *P. aeruginosa* to tobramycin in the community. We observed biofilm-specific tolerance of a Δ*lasR* mutant grown in the mixed community (**Fig. 3A**), a result repeated with two phenotypically LasR-defective CF clinical isolate variants (AMT0101-1-1; **Fig. 3C** and SMC1595; **Fig. S6**). Given that not all *P. aeruginosa* strains exhibited increased sensitivity in the mixed community, some strains may carry mutations in other pathways conferring tolerance to tobramycin in the mixed community. Indeed, clinical *P. aeruginosa* isolates can accumulate mutations including in the negative regulator MexZ, driving multidrug resilience (50).

Phenazines mediate the community tolerance of the Δ*lasR* mutant and can induce tolerance of WT *P. aeruginosa* (**Fig. 4B,C**). Our rationale for testing the impact of these redox-active compounds in our system is supported by reports highlighting the capacity of these molecules to drive *P. aeruginosa* drug tolerance (63-65). Also, strains lacking LasR function are known to overproduce phenazines (53, 58, 66-68). Interestingly, the highest levels of phenazine production by the Δ*lasR* mutant were observed in the mixed community (**Fig. 4A**). Our findings suggests that community members might impact phenazine production by *P. aeruginosa* grown in a polymicrobial environment; uncovering the mechanism of community-mediated stimulation of phenazine production is the goal of ongoing studies.

While the approaches used here have allowed us to explore community function, we acknowledge certain limitations to this study. First, while we provide a rationale for the selection of *P. aeruginosa, S. aureus, S. sanguinis*. and *P. melaninogenica* for use in our mixed community system, it is fair to ask, “is four enough?”. While the four microorganisms we have selected are a reasonable start, the model could be further improved by including additional CF-relevant species. However, adding pathogens such as *Burkholderia* or *Achromobacter* does not markedly increase the number of pwCF captured by our model (**Fig. S1B**,**C**). Furthermore, while the *in vitro* polymicrobial model does not perfectly match the Pa.M1 and Pa.M2 mixed community compositions detected in the CF airway (21), the abundances of *P. aeruginosa, S. aureus, S. sanguinis* and *P. melaninogenica* all fall within the clinical range detected in these communities (**Fig. S2A**,**B**). More importantly, the increased sensitivity of *P. aeruginosa* to tobramycin treatment in the mixed community could be observed for *P. aeruginosa* co-cultivated over a 1000-fold range of the other microbial partners (**Fig. S5**). These *in vitro* observations confirm the importance of studying microbes in the context of mixed-species communities. Finally, interrogating host factors (immune cells, host-derived metabolites) will also be an important next step.

## MATERIALS AND METHODS

Materials and methods for this study, including bacterial strains and culture conditions, microbial assays on plastic, drug susceptibility assays, minimal bactericidal concentration assays, time courses, high performance liquid chromatography-mass spectrometry (HPLC-MS/MS) quantification of molecules, and bioinformatics analyses are described in detail in the SI Appendix, SI Material and Methods section.

## Acknowledgements

This work was supported by NIH grant R01 AI155424 to G.A.O., Canadian Institutes of Health Research (CIHR) operating grant MOP-142466 to E.D., NIH/NIGMS 5 P20 GM130454 project support to D.S., Cystic Fibrosis Foundation (CFF) grant HOGAN19G0 to D.A.H. and CFF Postdoctoral Fellowship JEAN21F0 to F.J.P.. Additional support is provided by the Cystic Fibrosis Foundation Research Development Program (STANTO19R0) and NIH P30-DK117469 (Dartmouth Cystic Fibrosis Research Center). We thank Dallas L. Mould for technical help and helpful discussions. We also thank Sophie Robitaille, Courtney E. Price and Alexis R. Ramsey for helpful discussions.

