## Supplemental Data for "Community composition shapes microbial-specific phenotypes in a cystic fibrosis polymicrobial model system"

PNAS

**This PDF file includes:**

Supplementary Materials and Methods
Figures S1 to S15
Tables S1 to S2
SI References

**Supplemental Materials and Methods.**

**Bacterial strains and culture conditions.** All the *Pseudomonas aeruginosa*, *Staphylococcus aureus*, *Streptococcus* spp. and *Prevotella* spp. strains used in this study are listed in **Table S2**. *P. aeruginosa* and *S. aureus* cultures were grown in Tryptic Soy Broth (TSB) with shaking at 37°C. *Prevotella* spp. cultures were grown in TSB supplemented with 0.5% Yeast Extract (YE), 5 µg/mL hemin, 2.85 mM L-cysteine hydrochloride and 1 µg/mL menadione (*Prevotella* growth medium – PGM). *Streptococcus* spp. cultures were grown in Todd-Hewitt Broth supplemented with 0.5% YE (THY) at 37°C with 5% $CO_2$. Single colonies of each microbial species cultivated on either TSB solidified with 1.5% agar (TSA) for both *P. aeruginosa* and *S. aureus* species, or TSA supplemented with 5% sheep's blood (blood agar) for *Streptococcus* spp. and *Prevotella* spp. were utilized to start overnight cultures in the abovementioned liquid media. Artificial sputum medium with 5 mg/mL of mucin (ASM) was prepared as previously described (1, 2) and supplemented with 100 mM 3-morpholinopropane-1-sulfonic acid (MOPS) to maintain a pH of 6.80 over the course of our studies.

**Microbial assays.** Cultures experiments were performed using polystyrene flat-bottom 96-well plates. Cells from overnight liquid cultures of *P. aeruginosa*, *S. aureus*, *Streptococcus* spp. and *Prevotella* spp. were individually collected and washed twice (for *P. aeruginosa* and *S. aureus*) or once (for *Streptococcus* spp. and *Prevotella* spp.) in sterile Phosphate-Buffered Saline (PBS) by centrifuging at 10,000 x *g* for 2 minutes. After the final wash, cells were resuspended in ASM with no mucin. The optical density ($OD_{600}$) was then measured for each bacterial suspension and diluted to an $OD_{600}$ of 0.2 in ASM. Monocultures and co-culture conditions were prepared from the $OD_{600}$ = 0.2 suspension and diluted to a final $OD_{600}$ of 0.01 for each microbial species in ASM. A volume of 100 µl of bacterial suspension added to three wells. Plates were incubated using an AnaeroPak-Anaerobic container with a GasPak sachet (ThermoFisher) at 37°C for 24 hours. Then, unattached cells were aspirated with a multichannel pipette and the pre-formed biofilms replenished with 100 µl of fresh ASM and incubated for an additional 24 hours at 37°C, under anoxic conditions. After a total of 48 hours of incubation, the planktonic fraction was collected, 10-fold serially diluted and plated on *Pseudomonas* Isolation Agar (PIA) for *P. aeruginosa* detection, Mannitol Salt Agar (MSA) for *S. aureus* quantification, TSB + 0.5% YE + 1.5% agar supplemented with 5% sheep's blood, 10 µg/mL oxolinic acid, 10 µg/mL polymixin B for *Streptococcus* spp. (*Streptococcus* Selective Agar – SSA), TSB + 0.5% YE + 1.5% agar supplemented with 5% sheep's blood, 5 µg/mL hemin, 2.85 mM L-cysteine hydrochloride, 1 µg/mL menadione, 5 µg/mL vancomycin, 100 µg/mL kanamycin for *Prevotella* spp. (*Prevotella* Selective Agar – PSA). For surface-attached communities, biofilms were washed twice with sterile PBS and resuspended in 50 µL sterile PBS. Cells were disrupted using a 96-pin metal replicator and 10-fold serially diluted in sterile PBS. Dilutions were plated on PIA, MSA, SSA and PSA selective plates. *P. aeruginosa* and *S. aureus* selective plates were incubated for 16-20 hours at 37°C. *Streptococcus* spp. and *Prevotella* spp. selective plates were incubated in anoxic conditions for 24-48 hours at 37°C. After incubation, colonies were counted, and CFU/mL were determined for each microbial species. For the time course assays, CFU counts were performed for planktonic and biofilm fractions at 0, 3, 6, 12, 24, 48, 72, 96, 120, 144, 168 and 336 hours of monoculture and mixed communities grown in anoxic conditions. Pre-formed biofilms were replenished with fresh ASM every 24 hours for the duration of the experiment. For experiments with varying concentrations of *S. aureus*, *Streptococcus sanguinis* and *Prevotella melaninogenica* in monocultures and co-cultures, these species were grown from bacterial suspensions adjusted to an $OD_{600}$ = 0.8 in ASM. Suspensions were further diluted in ASM to an $OD_{600}$ of either 0.1, 0.001, 0.0001 or 0.00001 while maintaining *P. aeruginosa* at $OD_{600}$ = 0.01 in all conditions.

**Tobramycin susceptibility assay.** Fresh stocks of tobramycin sulfate (Alfa Aesar) were prepared in sterile milli-Q water to a concentration of 50 mg/mL and subsequently diluted to a working stock of 5 mg/mL in ASM for each experiment. Triplicate wells containing pre-formed biofilms of *P. aeruginosa*, *S. aureus*, *Streptococcus* spp. and *Prevotella* spp. grown as monoculture and mixed communities were initially grown for 24 hours as described in the "Microbial assays" section. After incubation, the non-attached cells were collected, and pre-formed biofilms exposed to 100 µg/mL tobramycin (100 µL/well) and further incubated for an additional 24 hours under anoxic conditions at 37°C. After a total of 48 hours of incubation, planktonic and biofilm cell fractions were sampled, ten-fold serially diluted and plated as described previously.

**Minimal bactericidal concentration (MBC) assay**. Communities were grown in ASM for a total of 48 hours as described in the “Microbial assays” section. The supernatants were collected then centrifuged for 10 minutes at 10,000 x *g* and filtered-sterilized by using at 0.22 µM filter. To assess sterility of the supernatants, a volume of 5 µL for each condition was spotted on PIA, MSA, SSA, PSA and blood agar plates. The supernatant spots were incubated at 37°C for PIA, MSA and blood agar plates (aerobically) as well as SSA, PSA and blood agar plates (anaerobically) for 24 hours. Serial two-fold dilutions of tobramycin ranging from 500 µg/mL to 0.49 µg/mL were performed in either ASM or with the supernatants from *P. aeruginosa* monocultures and mixed communities. Duplicate wells of a sterile 96-well plate were used to inoculate 50 µl of various tobramycin concentration ranges in combination with either (i) 50 µl of sterile ASM (negative control) or (ii) 50 µl of the monoculture or mixed community supernatants. The plate was incubated at room temperature for a total of 24 hours. To compare the MBC concentrations of tobramycin after incubation in ASM or in supernatants, overnight cultures of *P. aeruginosa* PA14 were washed twice with sterile 1X PBS and resuspended in ASM to a final $OD_{600}$ of 0.1. Fifty microliters of the *P. aeruginosa* bacterial suspension were then transferred to two wells of the 96-well plate containing the various two-fold dilutions of tobramycin made in either ASM or the culture supernatants. The plate was then incubated in aerobic conditions for 24 hours at 37°C. After the incubation, a 96-pin metal replicator was used to disrupt the cells formed at the bottom of the wells and then 3 µl of the resuspended cells were spotted on TSA plates and incubated at 37°C for 24 hours. Then, the MBC was determined by looking for absence of growth on the agar plate.

**Quantification of phenazines produced by *P. aeruginosa* by liquid chromatography tandem mass spectrometry (LC-MS/MS).** For phenazines quantification, communities of *P. aeruginosa* grown as monocultures and in mixed communities were prepared as described in the “Microbial assays” section and inoculated in twelve wells (full row) of a 96-well plate. After incubation under anaerobic conditions at 37°C, biofilms were disrupted into the planktonic fraction using at 96-well metal pin replicator, and cell suspension of ten wells were transferred to a sterile 2.0 mL microtube. A volume of 1 mL of 100% dichloromethane (DCM - Thermo Scientific Chemicals) containing 0.02 ppm of 5,6,7,8-tetradeutero-4-hydroxy-2-heptylquinoline (HHQ-d4) as the internal standard (IS) was then added to the cells. The samples were then vortexed for 20 seconds and centrifuged at 16,000 x *g* for 10 minutes at 4°C. The organic (bottom) phase of each sample was then transferred to a new 2.0 mL microtube. A second extraction of the remaining aqueous (top) phase was performed and added to the 1st organic extraction. The samples were evaporated at room temperature to a volume of 500 µL and then transferred to a conical HPLC glass vial before being completely evaporated. Samples were stored at -20°C until resuspended in 100 µL HPLC-grade acetonitrile and analysed using liquid chromatography tandem mass spectrometry (LC-MS/MS) as previously described with minor modifications (3). A 15 µL volume of sample was injected and analyzed by HPLC (Waters 2795; Waters, Mississauga, ON, Canada) using a Kinetex (100- by 3.0-mm) 5-µm EVO $C_{18}$ reverse-phase LC column (Phenomenex). The detector was a tandem quadrupole mass spectrometer (Quattro premier XE; Waters) equipped with a Z-spray interface using electrospray ionization in positive mode (ESI+). Nitrogen was used as a nebulizing and drying gas at flow rates of 15 and 100 ml · $min^{-1}$, respectively. HPLC flow rate was 400µl/min split to 40 µl/min by a Valco tee splitter. An acetonitrile-water gradient containing 1% acetic acid was used. In MRM mode, the following transition were monitored: 225 → 207 for phenazine-1-carboxylic acid (PCA) and 248→163 for the internal standard. The collision energies were set at 15, and 30 V respectively and the collision gas flow (argon) was set at 0.35 ml/min. Pure PCA (Sigma) was used as a standard for the method.

**Quantification of 3-oxo-$C_{12}$ homoserine lactone produced by *P. aeruginosa* by liquid chromatography tandem mass spectrometry (LC-MS/MS).** Extraction of the LasR regulated signaling molecule 3-oxo-$C_{12}$ homoserine lactone (3-oxo-$C_{12}$-HSL) from pure culture bacterial suspensions was performed as described in the "Quantification of phenazines produced by *P.*

*aeruginosa* by liquid chromatography tandem mass spectrometry (LC-MS/MS)" section. Quantification of 3-oxo-$C_{12}$-HSL was done as previously reported by (3).

**Tobramycin susceptibility assay in the presence of phenazines.** Fresh stocks of tobramycin sulfate (Alfa Aesar) were prepared in sterile milli-Q water to a concentration of 50 mg/mL and subsequently diluted to a working stock of 5 mg/mL in ASM for each experiment. A 20 mM pyocyanin (Sigma) stock was prepared in 100% DMSO. Preparations containing either (i) ASM with or without vehicle controls, (ii) ASM supplemented with 100 µg/mL of tobramycin, (iii) ASM supplemented with 50 µM or 300 µM pyocyanin and (iv) ASM supplemented with 100 µg/mL tobramycin and 50 µM or 300 µM pyocyanin, were added to triplicate wells containing pre-formed biofilms of *P. aeruginosa*, *S. aureus*, *S. sanguinis* and *P. melaninogenica* grown as monoculture and mixed community biofilms grown for 24 hours as described in the "Microbial assays" section. Communities were incubated for 24 hours in anoxic conditions at 37°C. After a total of 48 hours of incubation, biofilm cell fractions were taken, ten-fold serially diluted and plated as described previously.

**Molecular techniques**. In frame deletion of the *lasR* gene in a ΔΔ*phz* mutant was done using *E. coli* SM10 λpir carrying the pEX18Gm-Δ*lasR* plasmid as previously reported (4). Complementation of the *lasR* gene in a Δ*lasR* mutant background was done using *E. coli* S17 λpir carrying the knock-in pMQ30-*lasR* construct as previously described (5). Strains were confirmed by sequencing.

**16S rRNA gene data analyses**. The compendium of CF sputum microbiome data sets used here was developed from publicly available 16S rRNA reads as previously published (6). Briefly, reads from sputum samples from 167 clinically stable subjects aged 8 to 69 from 14 CF centers in the United States and Europe were assigned to bacterial taxa at the genus level. Applying a gap statistic to unsupervised k-means clustering of Bray-Curtis distance of these 167 subject samples suggested that 5 community types best represented differences between the 167 subjects as previously described (6). Here we have used this compendium of taxonomically assigned reads from 167 pwCF to assess how many distinct genera, which ones, are required to capture 70% or more of taxonomically assigned reads in each subject.

***In silico* community modeling**. We used the SteadyCom method (7) to simulate the steady-state metabolism of a *P. aeruginosa*, *S. aureus*, *S. sanguinis* and *P. melaninogenica* mixed community. SteadyCom performs community flux balance analysis by computing the maximal community growth while ensuring that all metabolites are properly balanced within each species and across the community. SteadyCom provides the capability to constrain the relative abundance of each species in the community to determine the main interspecies metabolic interactions. This simulation method is based on several simplifying assumptions, including spatial homogeneity and that all propagating species have the same growth rate at steady state. Outputs of each SteadyCom simulation included the community growth rate and species-dependent uptake and secretion rates of each extracellular metabolite. Model parameters, such as constraints in metabolite uptake rates, were taken from previous work modeling cystic fibrosis airway communities (8). Four independent simulations, corresponding to the species abundances in the community types found to explain the variability in lung function were ran. We calculated the relative abundance between *P. aeruginosa*, *S. sanguinis*, *P. melaninogenica* and *S. aureus* within each community type, while ignoring remaining species. The simulations were then used to determine the main metabolites exchanged between the four species.

Species abundance in each community type (based from (6))

| | *P. aeruginosa* | *S. sanguinis* | *P. melaninogenica* | *S. aureus* |
|---|---|---|---|---|
| *Pseudomonas*-dominated (Pa.D) | 77.6% | 3.0% | 6.2% | 1.6% |
| *Streptococcus*-dominated (Strep.D) | 4.4% | 49.5% | 16.9% | 1.7% |
| Pa.M1 | 55.6% | 13.6% | 10.2% | 1.2% |
| Pa.M2 | 67.7% | 2.7% | 10.4% | 2.5% |

**Protease assays**. Protease activity was assessed through casein degradation of tested *P. aeruginosa* strains grown on 1.5% TSA plates containing 1.5% sterile skim milk. Briefly, six colonies and appropriate controls grown on TSA plates were inoculated on milk plates and incubated overnight at 30°C for 24 and 48 hours. Casein degradation activity was observed by the presence or not of a zone of clearance around the inoculation site.

**3-oxo-$C_{12}$-HSL autoinducer assays.** The production 3-oxo-$C_{12}$-HSL of tested strain and associated controls was measured as described previously through the utilization of a 33-oxo-$C_{12}$-HSL specific *lacZ* reporter (9). Briefly, the 3-oxo-$C_{12}$-HSL-specific *lacZ* reporter strain (DH161) was diluted to an $OD_{600}$ = 0.01 in LB from an overnight culture. A volume of 100 µl was spread on LB agar plates supplemented with 150 µg/mL of 5-bromo-4-chloro-3-indolyl-β-D-galactopyranoside (X-gal) dissolved in 100% DMSO. Then, the plates were allowed to dry for 15 minutes in a biological safety cabinet. Once dry, a volume of 5 µL of $OD_{600}$-adjuted to 1.0 tested strains and controls (PA14 wild type, Δ*lasR* and Δ*lasI*Δ*rhlI*) was spotted on the pre-inoculated reporter strain. The plates were allowed to dry for an additional 15 minutes and then incubated at 37°C for 16-18 h. Development of a blue halo around tested colonies was interpreted as a positive 3-oxo-$C_{12}$-HSL activity.

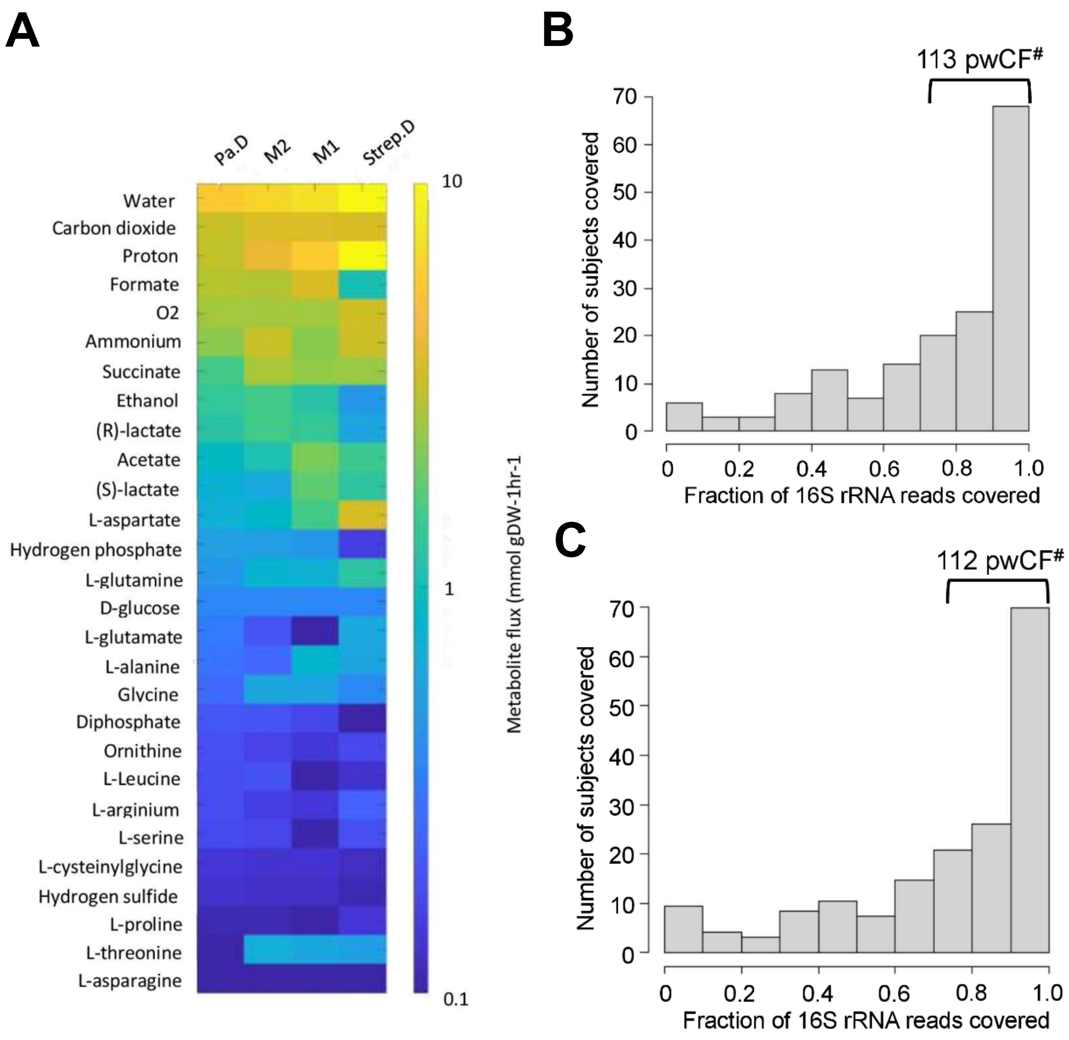

**Fig. S1. Leveraging clinical microbiome data sets with computational analyses to identify communities and community members to model *in vitro*.** (**A**) *In silico* prediction of metabolic flux between *P. aeruginosa*-dominated (Pa.D), *Streptococcus*-dominated (Strep.D) and the "mixed" Pa.M1 and Pa.M2 communities based on the abundance of *P. aeruginosa*, *S. aureus*, *S. sanguinis* and *P. melaninogenica* analyzed by Hampton, O'Toole and colleagues (6). (**B, C**) Number of unique samples for which $\geq$70% of 16S rRNA reads can be associated with the combined presence of *Pseudomonas*, *Staphylococcus*, *Streptococcus*, *Prevotella* plus either *Burkholderia* (**B**) or *Achromobacter* (**C**). [#]Indicates the number of samples that meet these criteria from the total sample size is 167 persons with CF (pwCF). Compare the values in panels **B,C** with the 103 samples with >70% *Pseudomonas*, *Staphylococcus*, *Streptococcus*, *Prevotella* shown in **Figure 1B**.

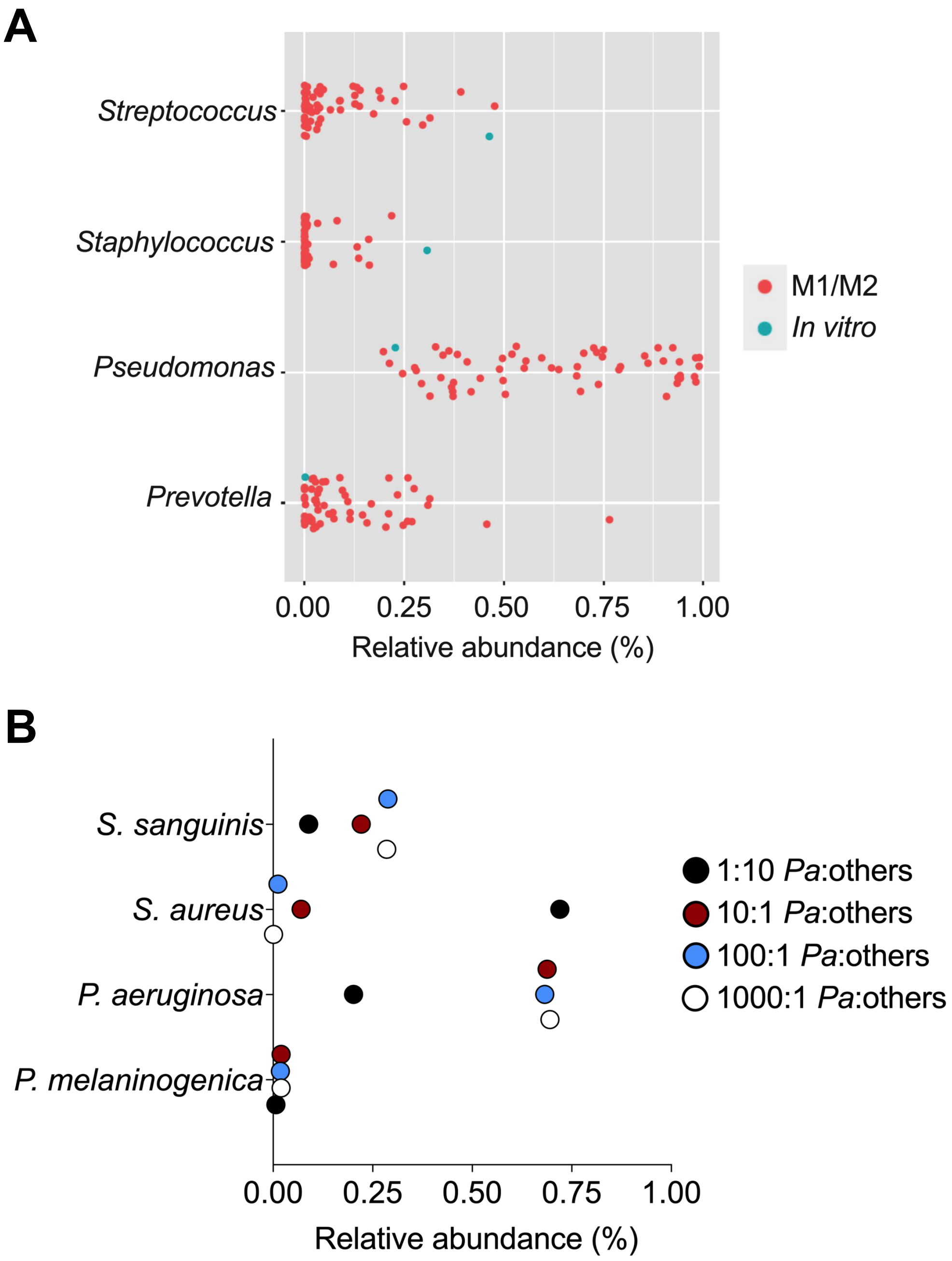

**Fig. S2. Microbial composition range of *in vivo* CF mixed communities compared with the model.** (**A**) Relative abundance ranges of 16S rRNA reads assigned to *Pseudomonas*, *Staphylococcus*, *Streptococcus* and *Prevotella* detected in the M1/M2 community types identified previously (6). These data were plotted against the relative abundance of *P. aeruginosa*, *S. aureus*, *S. sanguinis* and *P. melaninogenica* grown in the *in vitro* mixed community used in all the studies in this manuscript. (**B**) *In vitro* relative abundance of *P. aeruginosa*, *S. aureus*, *S. sanguinis* and *P. melaninogenica* in mixed communities where *P. aeruginosa* was maintained at the same starting inoculum and other microbes grown 10X in excess or diluted from 1:10 to 1:1000 relative to *P. aeruginosa*. *Pa* = *P. aeruginosa*, others = *S. aureus*, *S. sanguinis* and *P. melaninogenica*.

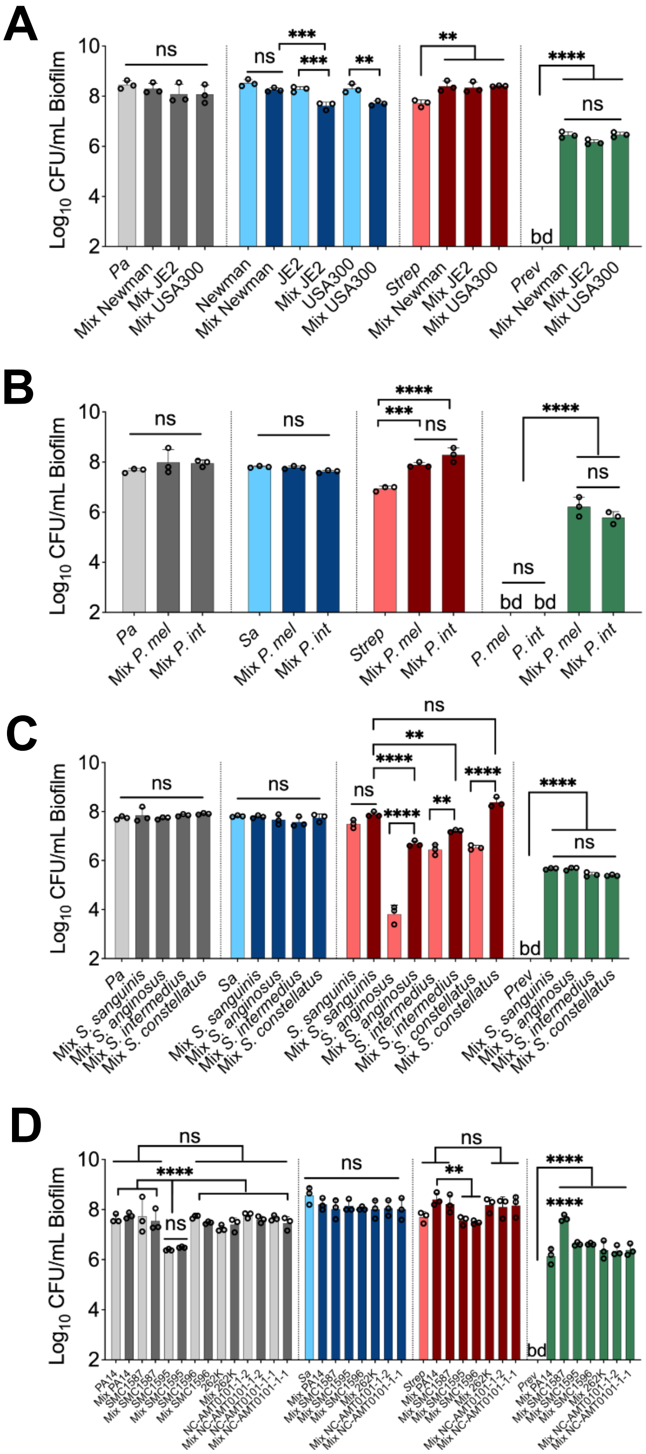

**Fig. S3. Testing additional laboratory and clinical strains in the *in vitro* polymicrobial community model**. Colony forming units (CFUs) of communities including (**A**) *S. aureus* strains, (**B**) *P. melaninogenica* (*P. mel*), *Prevotella intermedia* (*P. int*), (**C**) *Streptococcus* spp. and, (**D**) *P. aeruginosa* strains. Each column represents the average from at least three biological replicates, each with at least three technical replicates. Statistical analysis was performed using ordinary one-way analysis of variance (ANOVA) and Tukey's multiple comparisons posttest with *, $P < 0.05$; **, $P < 0.01$; ***, $P < 0.001$ and ****, $P < 0.0001$, ns = non-significant. *Pa* = *P. aeruginosa* PA14, *Sa* = *S. aureus* Newman, *Strep* = *S. sanguinis*, *Prev* = *P. melaninogenica*, bd = below detection.

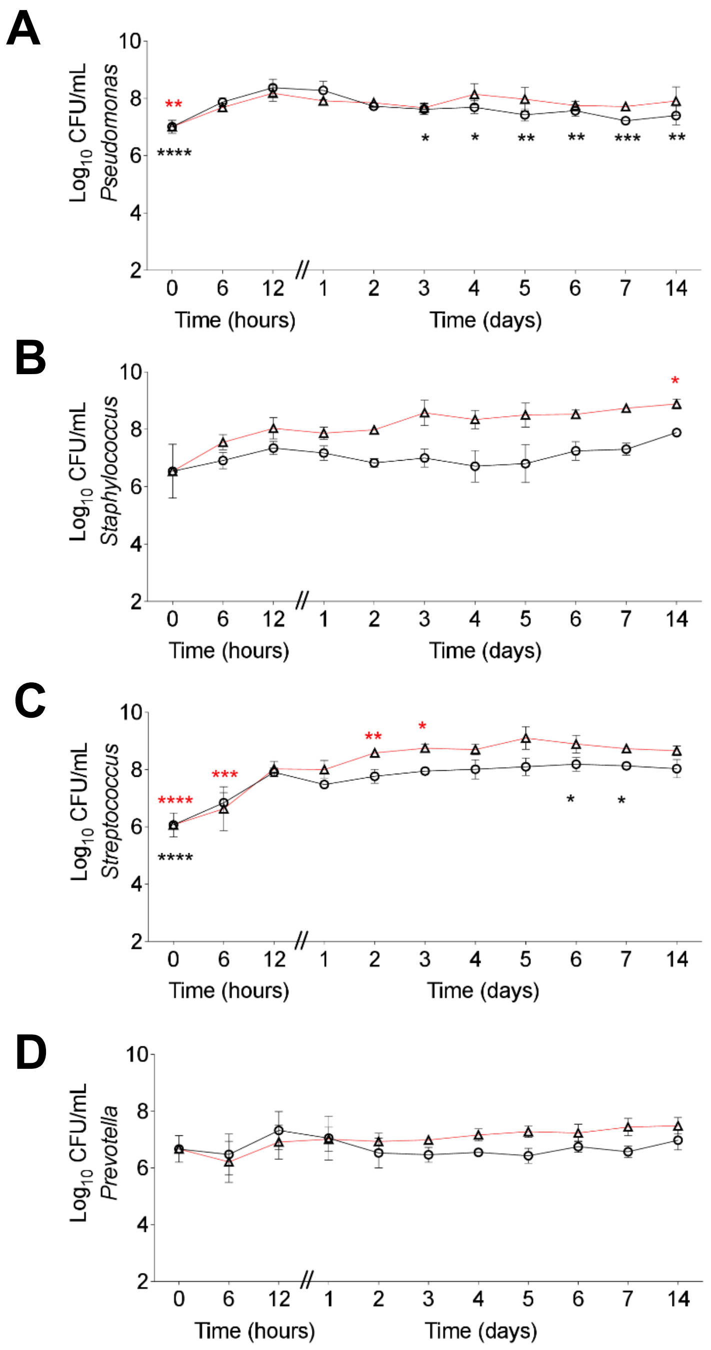

**Fig. S4. Fourteen day co-culture experiment of community members grown in a planktonic and biofilm mixed communities**. Colony forming units (CFUs) of (**A**) *P. aeruginosa*, (**B**) *S. aureus*, (**C**) *S. sanguinis* and (**D**) *P. melaninogenica*. Red lines represent biofilm cells and black lines, planktonic ones. Each timepoint represents the average from at least three biological replicates, each with at least three technical replicates. Every 24 hrs, the medium was removed, and then fresh medium added. Statistical analysis was done using ordinary one-way analysis of variance (ANOVA) and Dunnett's multiple comparisons posttest with the 24 hr timepoint as control. *, $P < 0.05$; **, $P < 0.01$; ***, $P < 0.001$; ****, $P < 0.0001$.

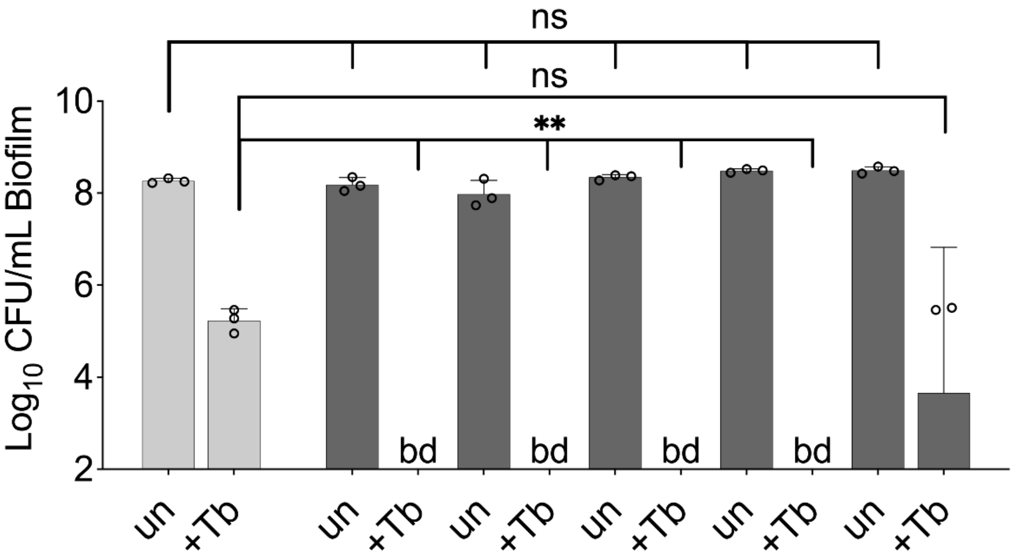

| <u>*Pa*:others</u> | | | | | | |
|---|---|---|---|---|---|---|
| *Pa* | 1 | 1 | 1 | 1 | 1 | 1 |
| *Sa* | - | 1 | 10 | 0.1 | 0.01 | 0.001 |
| *Strep* | - | 1 | 10 | 0.1 | 0.01 | 0.001 |
| *Prev* | - | 1 | 10 | 0.1 | 0.01 | 0.001 |

**Fig. S5. Microbial partners increase the killing of *P. aeruginosa* exposed to Tb in a mixed community over a wide range of population values.** Colony forming units (CFUs) of *P. aeruginosa* grown in the presence of varying concentrations of *S. aureus*, *S. sanguinis* and *P. melaninogenica* ranging from 10X the initial abundance of *P. aeruginosa* to 1000X less (0.0001) for all three of the other organisms, and then treated with tobramycin. *P. aeruginosa* was inoculated at the same starting $OD_{600}$ in all conditions. Each column represents the average from at least three biological replicates, each with at least three technical replicates. Statistical analysis was performed using ordinary one-way analysis of variance (ANOVA) and Tukey's multiple comparisons posttest with **, $P < 0.01$. ns = non-significant, bd = below detection, +Tb = + 100 µg/mL tobramycin, un = untreated. *Pa* = *P. aeruginosa*, *Sa* = *S. aureus*, *Strep* = *S. sanguinis*, *Prev* = *P. melaninogenica*. Monoculture biofilm = light grey, mixed biofilm = dark grey.

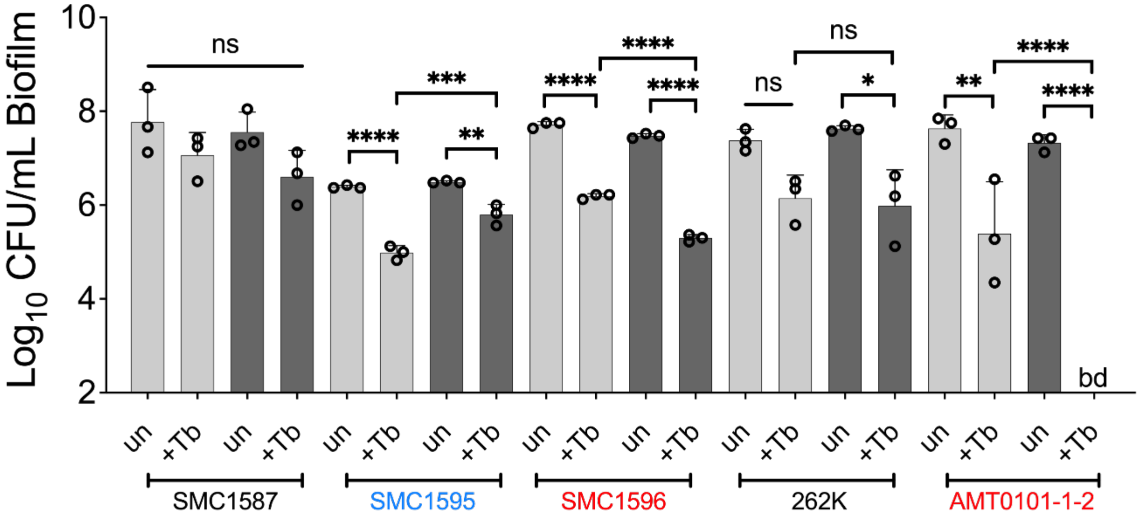

**Fig. S6. Drug sensitivity of *P. aeruginosa* clinical strains grown in a mixed community challenged with tobramycin**. Colony forming units (CFUs) of various *P. aeruginosa* strains grown as monocultures (light grey) or mixed biofilm communities (dark grey) treated with tobramycin. Each column represents the average from at least three biological replicates, each with at least three technical replicates. Statistical analysis was performed using ordinary one-way analysis of variance (ANOVA) and Tukey's multiple comparisons posttest with *, $P < 0.05$; **, $P < 0.01$; ***, $P < 0.001$ and ****, $P < 0.0001$, ns = non-significant, bd = below detection, +Tb = + 100 µg/mL tobramycin, un = untreated.

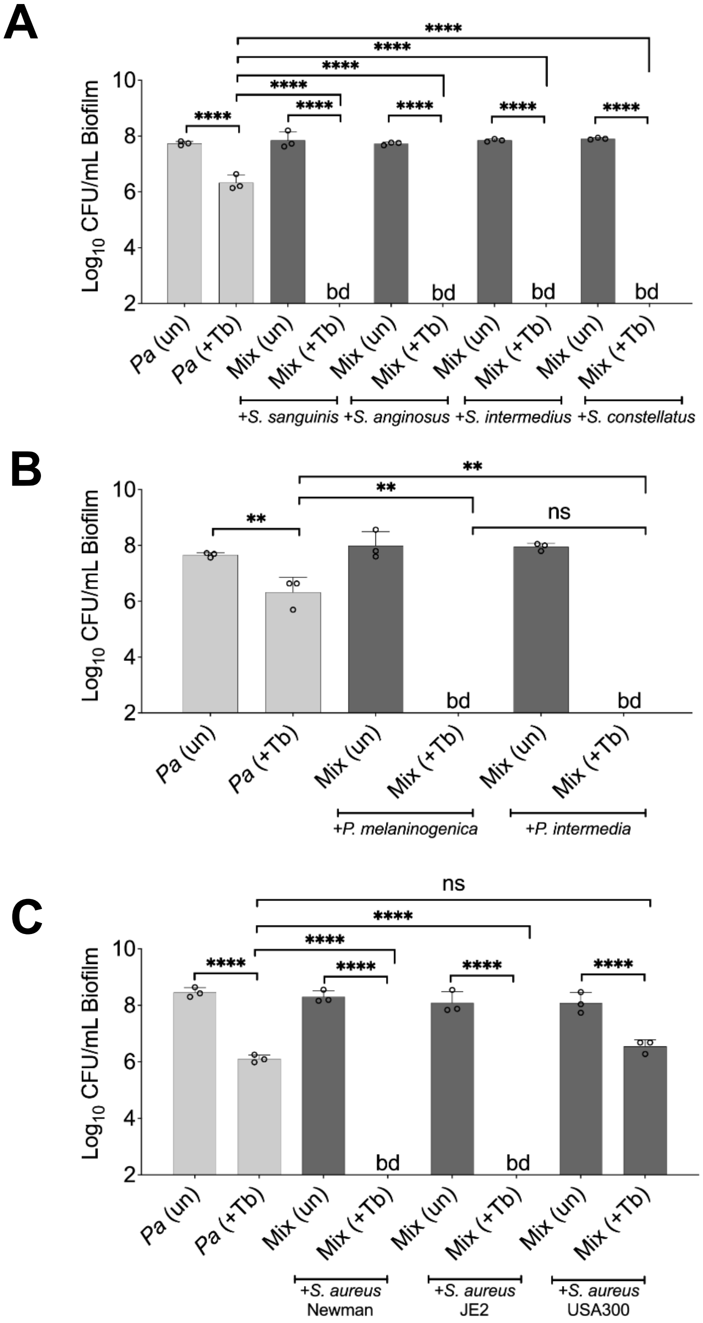

**Fig. S7. Shifted sensitivity of *P. aeruginosa* PA14 grown in a mixed community using various laboratory and clinical isolates treated with tobramycin.** Colony forming units (CFUs) of *P. aeruginosa* PA14 biofilms grown as monocultures (light grey) or mixed communities (dark grey) in the presence of (**A**) *S. sanguinis, Streptococcus anginosus, Streptococcus intermedius* and *Streptococcus constellatus*, (**B**) *P. melaninogenica*, *Prevotella intermedia* or, (**C**) *S. aureus* strains/species treated with tobramycin. Each column represents the average from at least three biological replicates, each with at least three technical replicates. Statistical analysis was performed using ordinary one-way analysis of variance (ANOVA) and Tukey's multiple comparisons posttest with **, $P < 0.01$ and ****, $P < 0.0001$, ns = non-significant. bd = below detection, *Pa* = *P. aeruginosa*, *P. mel* = *P. melaninogenica*; *P. int* = *Prevotella intermedia*, +Tb = + 100 µg/mL tobramycin, un = untreated.

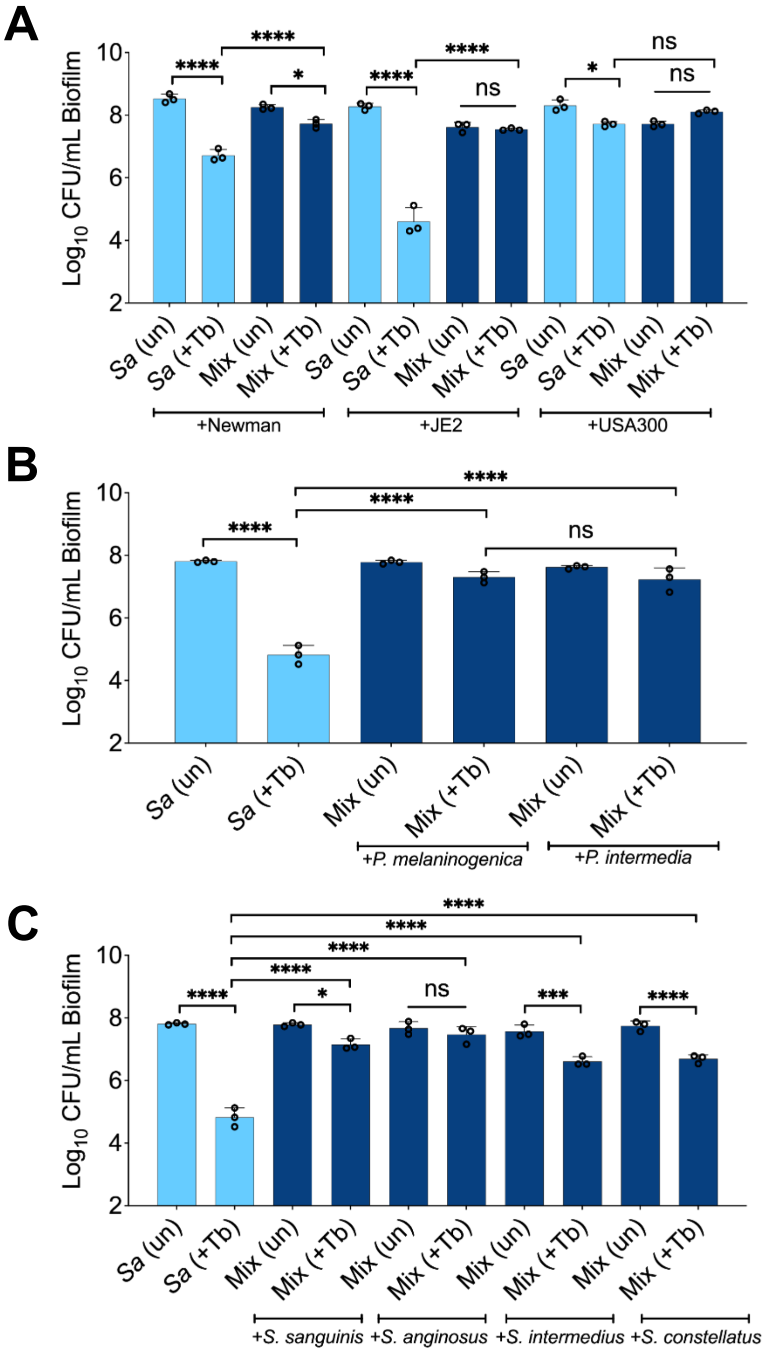

**Fig. S8. Recalcitrance of *S. aureus* biofilms grown in a mixed community comprised of various clinical isolates and treated with tobramycin**. *S. aureus* colony forming units (CFUs) when grown as monocultures (light blue) or mixed communities (dark blue) treated with tobramycin for (**A**) *S. aureus* strains spp., (**B**) *Prevotella* spp. and (**C**) *Streptococcus* spp. Each column represents the average from at least three biological replicates, each with at least three technical replicates. Statistical analysis was performed using ordinary one-way analysis of variance (ANOVA) and Tukey's multiple comparisons posttest with *, $P < 0.05$; ***, $P < 0.001$ and ****, $P < 0.0001$, ns = non-significant, bd = below detection, +Tb = +100 μg/mL tobramycin, un = untreated, *Sa* = *S. aureus* Newman, *P. inter* = *P. intermedia*, *P. mel* = *P. melaninogenica*.

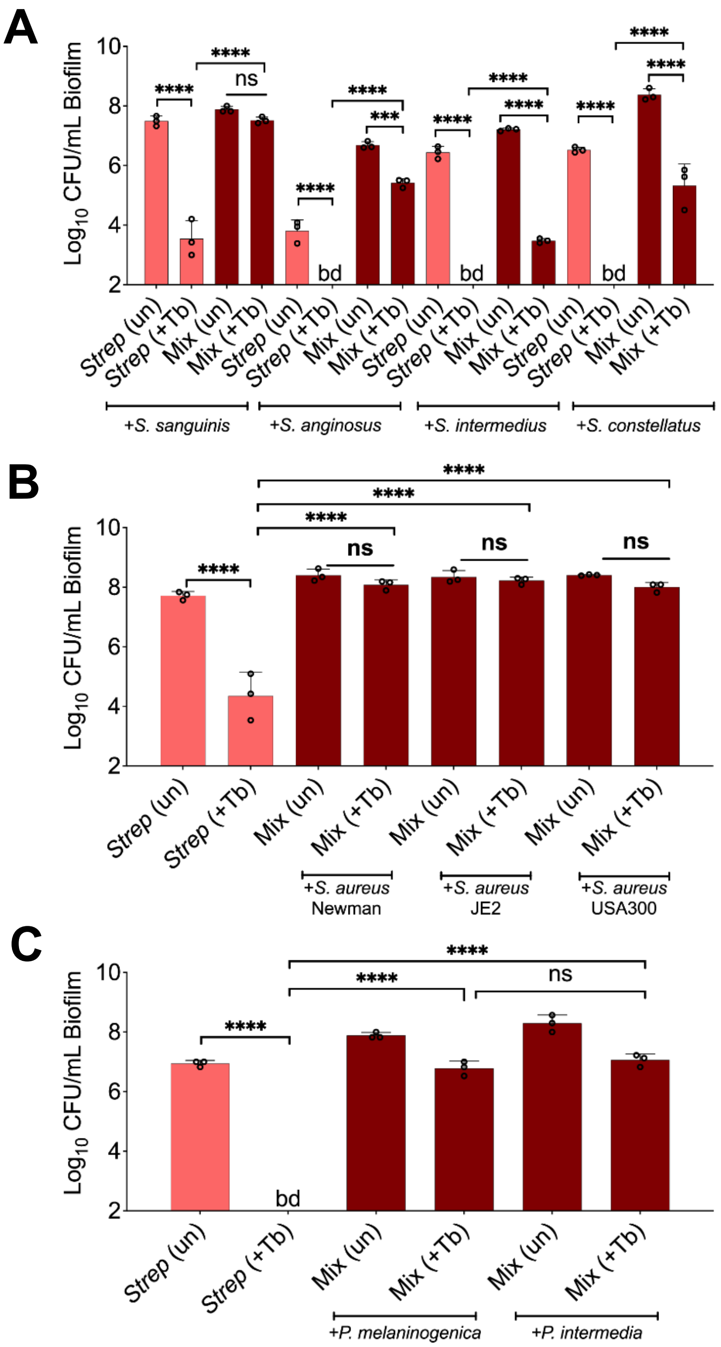

**Fig. S9. Recalcitrance of *Streptococcus* spp. biofilms grown in a mixed community comprised of various clinical isolates and treated with tobramycin**. *Streptococcus* spp. colony forming units (CFUs) when grown as monocultures (light red) or mixed communities (dark red) treated with tobramycin for (**A**) *Streptococcus* spp., (**B**) *S. aureus* strains or (**C**) *Prevotella* spp. Each column represents the average from at least three biological replicates, each with at least three technical replicates. Statistical analysis was performed using ordinary one-way analysis of variance (ANOVA) and Tukey's multiple comparisons posttest with ***, $P < 0.001$ and ****, $P < 0.0001$, ns = non-significant, bd = below detection, +Tb = +100 μg/mL tobramycin, un = untreated, *Strep* = *S. sanguinis*, *P. inter* = *P. intermedia*, *P. mel* = *P. melaninogenica*.

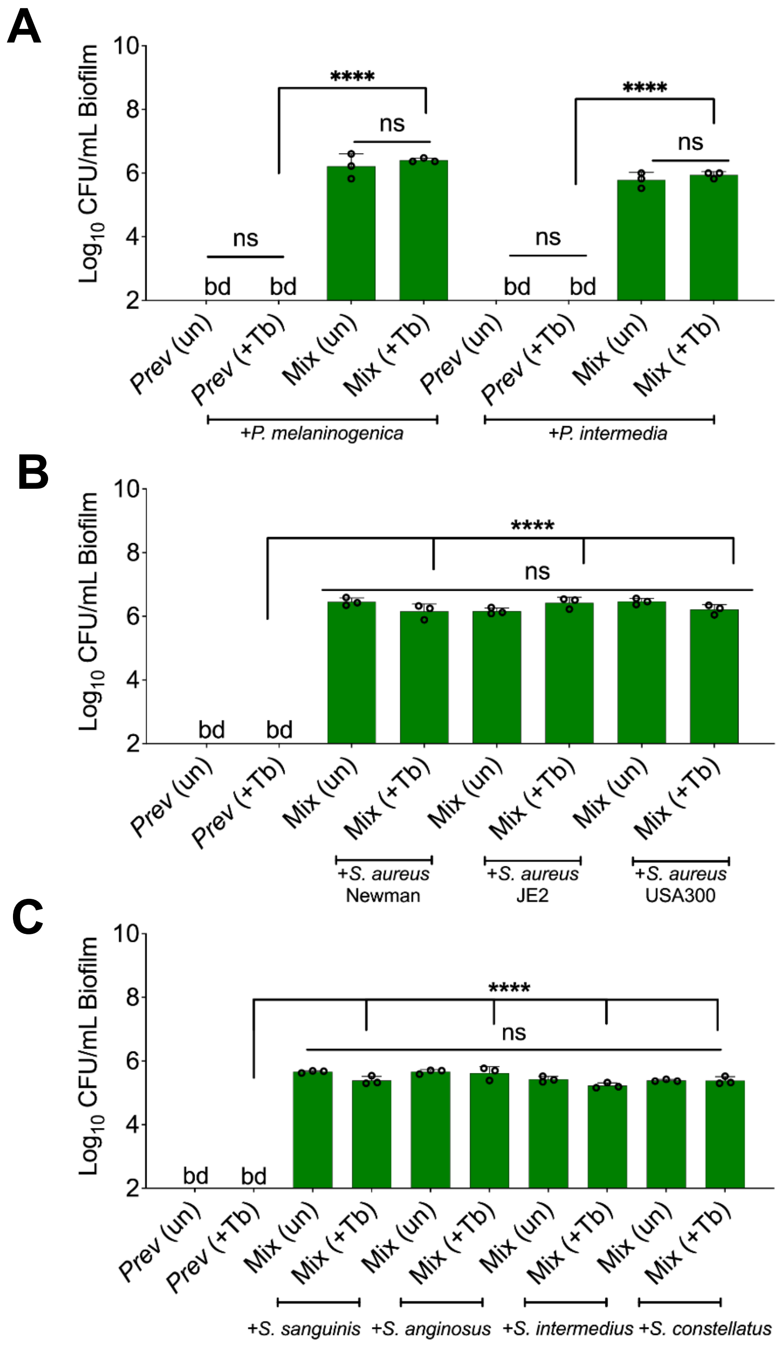

**Fig. S10**. **Recalcitrance of *Prevotella* spp. biofilms grown in a mixed community comprised of various clinical isolates and treated with tobramycin**. *Prevotella* spp. colony forming units (CFUs) when grown as monocultures (light green) or mixed communities (dark green) treated with tobramycin for (**A**) *Prevotella* spp., (**B**) *S. aureus* strains or (**C**) *Streptococcus* spp. Each column represents the average from at least three biological replicates, each with at least three technical replicates. Statistical analysis was performed using ordinary one-way analysis of variance (ANOVA) and Tukey's multiple comparisons posttest with ****, P < 0.0001, ns = non-significant, bd = below detection, +Tb = +100 μg/mL tobramycin, un = untreated, *P. inter* = *P. intermedia*, *P. mel* = *P. melaninogenica*.

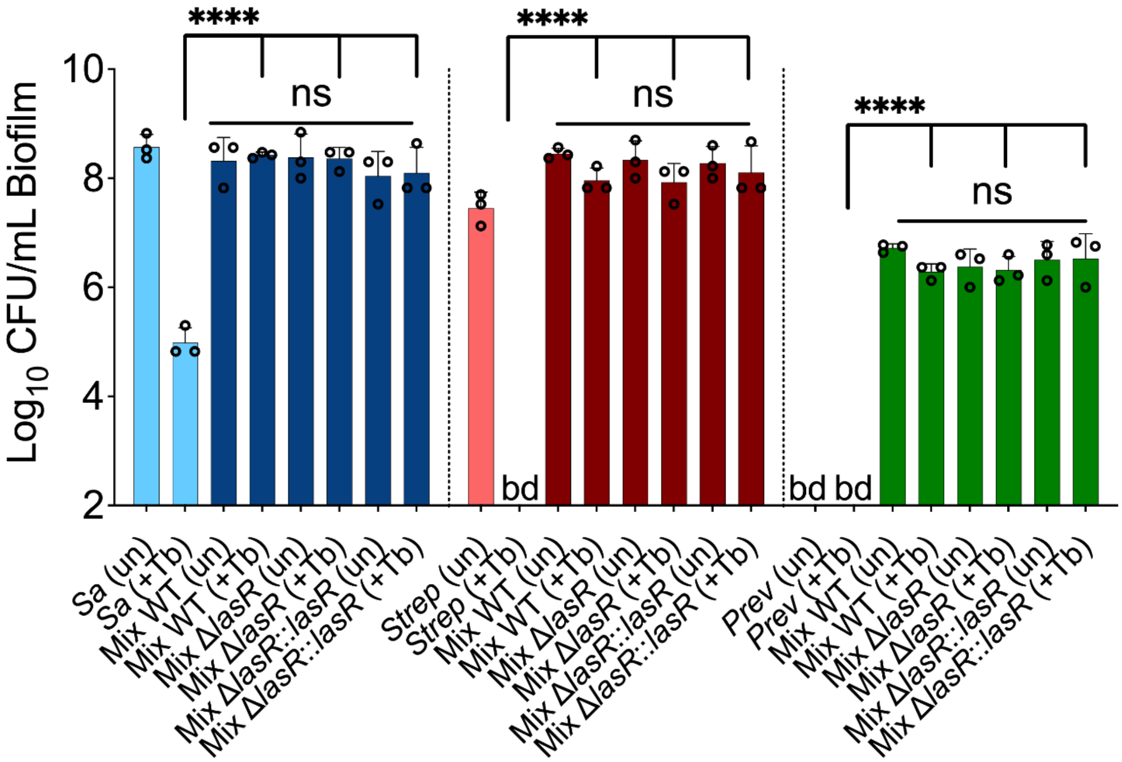

**Fig. S11. Loss of *P. aeruginosa* LasR function does not alter the viability of the other microbes in the mixed community compared to growth with WT *P. aeruginosa*.** Colony forming units (CFUs) counts of *S. aureus* (*Sa*), *S. sanguinis* (*Strep*) and *P. melaninogenica* (*Prev*) grown as monoculture (light color) or mixed biofilm communities (dark color) with WT *P. aeruginosa* (WT) and associated mutants treated with tobramycin. Each column represents the average from at least three biological replicates, each with at least three technical replicates. Statistical analysis was performed using ordinary one-way analysis of variance (ANOVA) and Tukey's multiple comparisons posttest with ****, $P$ < 0.0001, ns = non-significant, bd = below detection, un = untreated, +Tb = +100 µg/mL tobramycin.

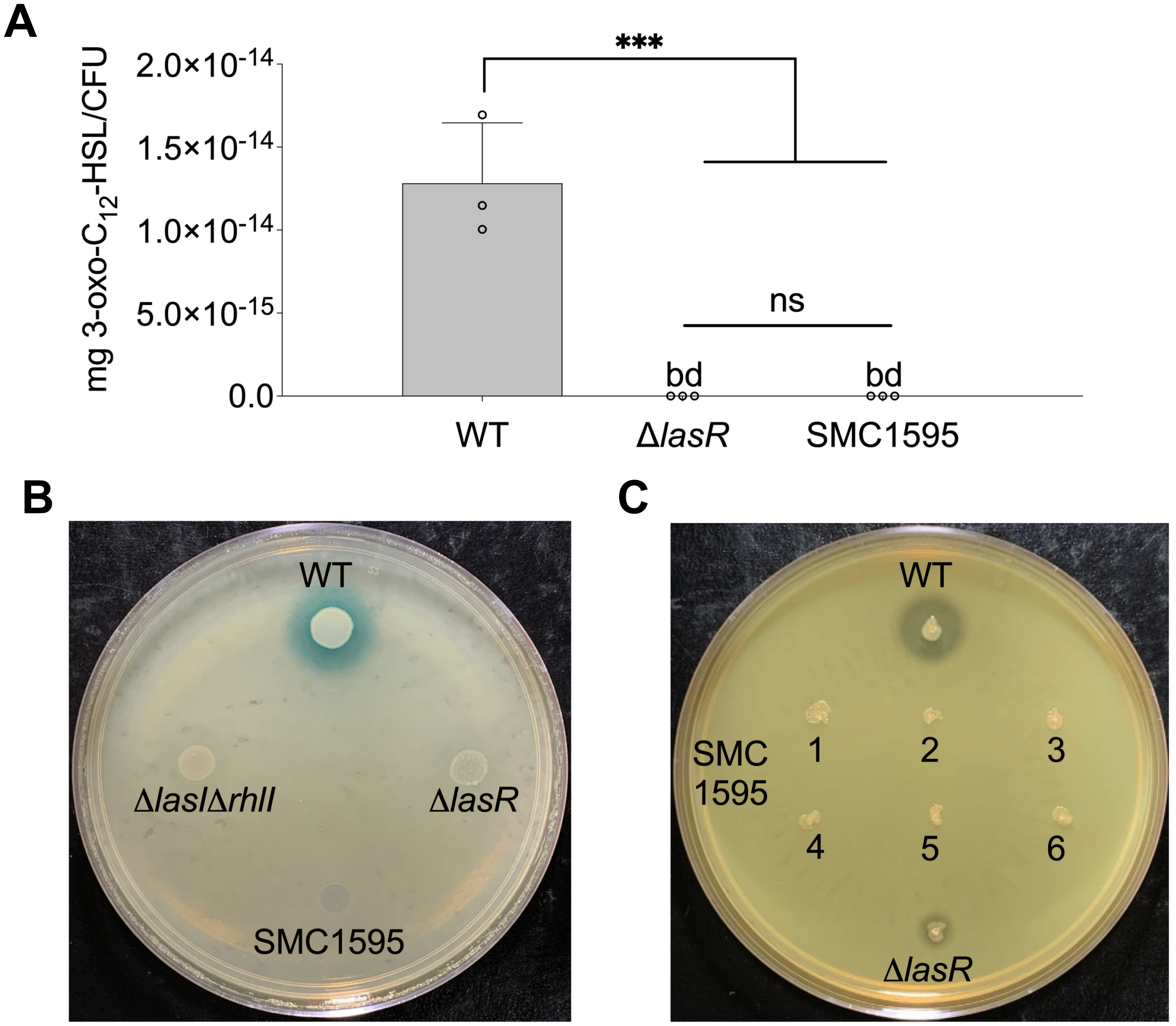

**Fig. S12. LasR-specific phenotypic tests of *P. aeruginosa* CF clinical isolate SMC1595.** (**A**) LC-MS/MS quantification of the 3-oxo-$C_{12}$-HSL signaling molecule. Each column represents the average from at least three biological replicates, each with at least three technical replicates. Statistical analysis was performed using ordinary one-way analysis of variance (ANOVA) and Tukey's multiple comparisons posttest with ***, $P < 0.001$, ns = non-significant, bd = below detection. (**B**) 3-oxo-$C_{12}$-HSL-specific *lacZ* bioreporter assay. (**C**) Protease activity on milk plates. For the bioreporter assays, four SMC1595 clones were tested at least on four different days. For protease assays, six clones of SMC1595 were tested at least on three different days. WT = *P. aeruginosa* PA14, ns = non-significant, bd = below detection.

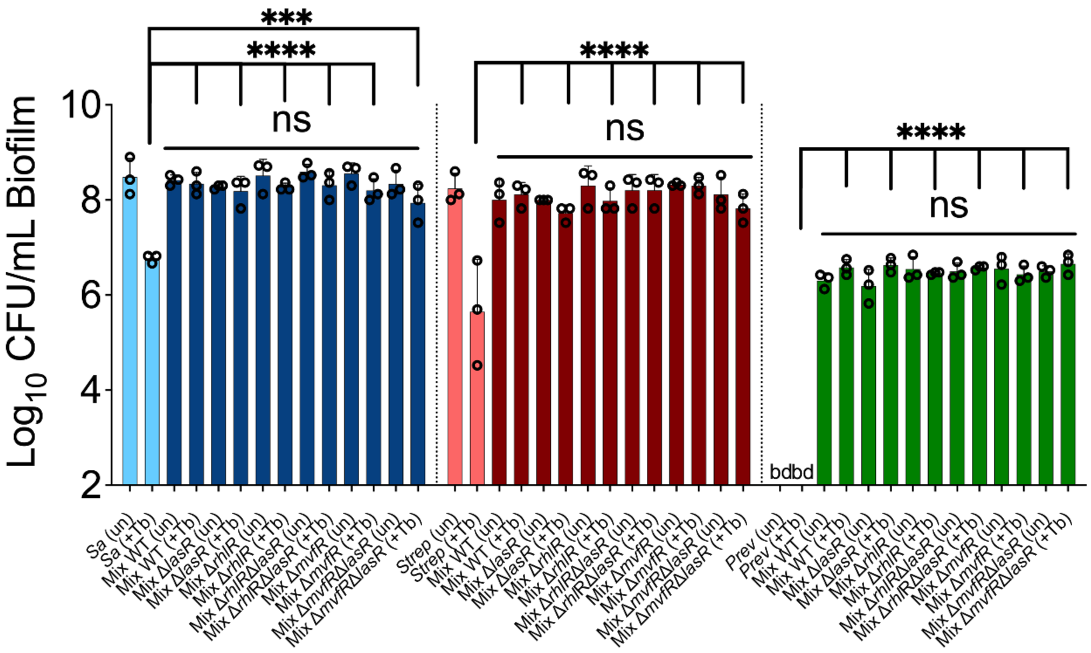

**Fig. S13. Tolerance of Δ*lasR* mutant in a mixed community: a role for the MvfR/PQS regulatory system.** Colony forming unit (CFU) counts of *S. aureus* (*Sa*), *S. sanguinis* (*Strep*) and *P. melaninogenica* (*Prev*) grown as monoculture (light color) or mixed biofilm communities (dark color) with WT *P. aeruginosa* (WT) or the indicated mutants treated with tobramycin. Each column represents the average from at least three biological replicates, each with at least three technical replicates. Each column represents the average from at least three biological replicates, each with at least three technical replicates. Statistical analysis was performed using ordinary one-way analysis of variance (ANOVA) and Tukey's multiple comparisons posttest with **, $P < 0.001$ and ****, $P < 0.0001$, ns = non-significant, bd = below detection, un = untreated, +Tb = +100 µg/mL tobramycin.

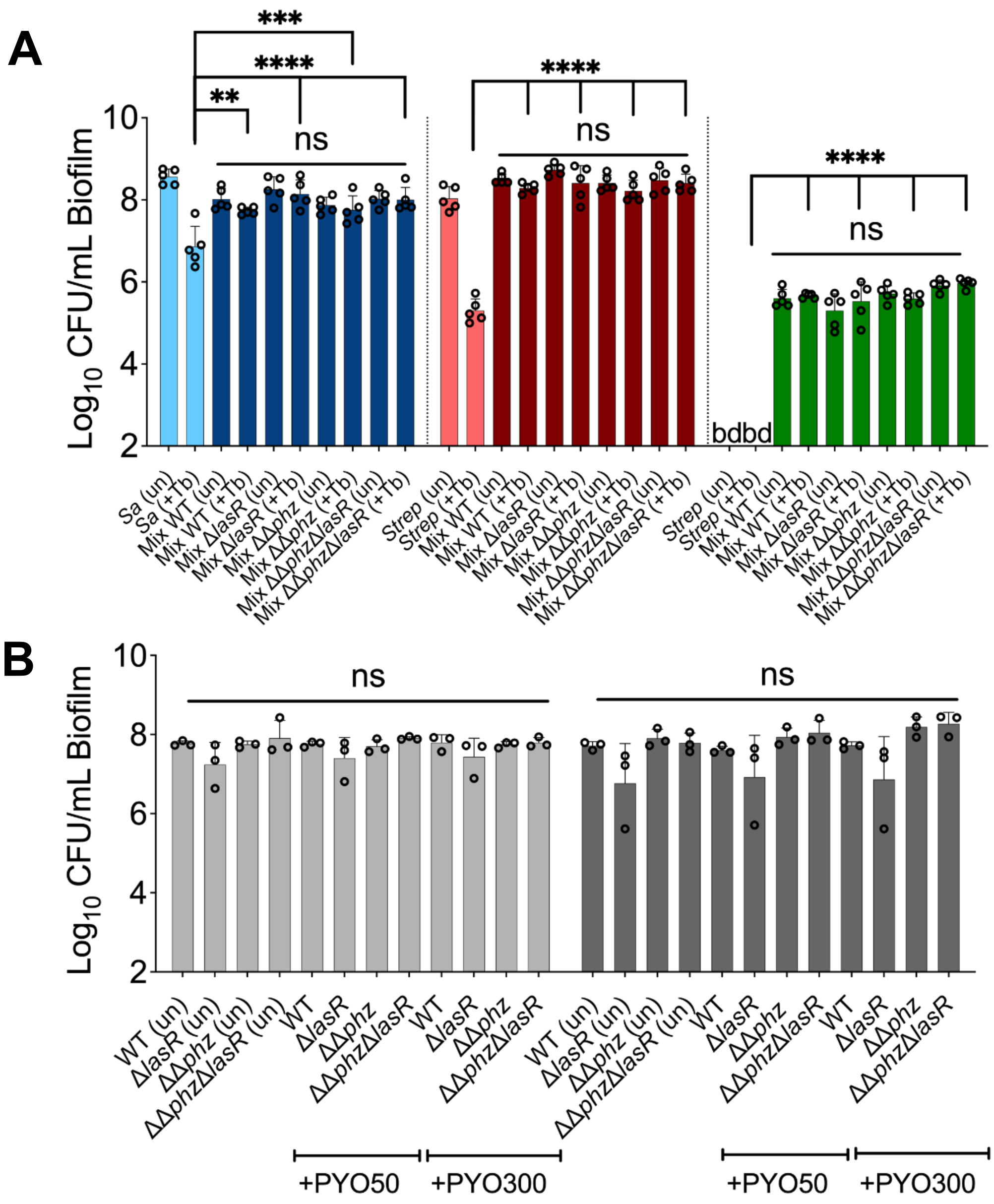

**Fig. S14. Phenazine drive tolerance of *P. aeruginosa* in mixed communities: lack of impact on other community members.** Colony forming units (CFUs) counts of (**A**) *S. aureus* (*Sa*), *S. sanguinis* (*Strep*) and *P. melaninogenica* (*Prev*) grown as monoculture (light color) or mixed biofilm communities (dark color) with WT *P. aeruginosa* (WT) and associated mutants treated with tobramycin. (**B**) CFU counts of monoculture (light grey), or mixed (dark grey) *P. aeruginosa* (WT) and associated mutants biofilm communities treated or not with 50 µM (+PYO50) and 300 µM (+PYO300) of phenazine. Each column represents the average from at least three biological replicates, each with at least three technical replicates. Statistical analysis was performed using ordinary one-way analysis of variance (ANOVA) and Tukey's multiple comparisons posttest with **, $P < 0.01$; ***, $P < 0.001$, and ****, $P < 0.0001$, ns = non-significant, bd = below detection, un = untreated, +Tb = +100 µg/mL tobramycin.

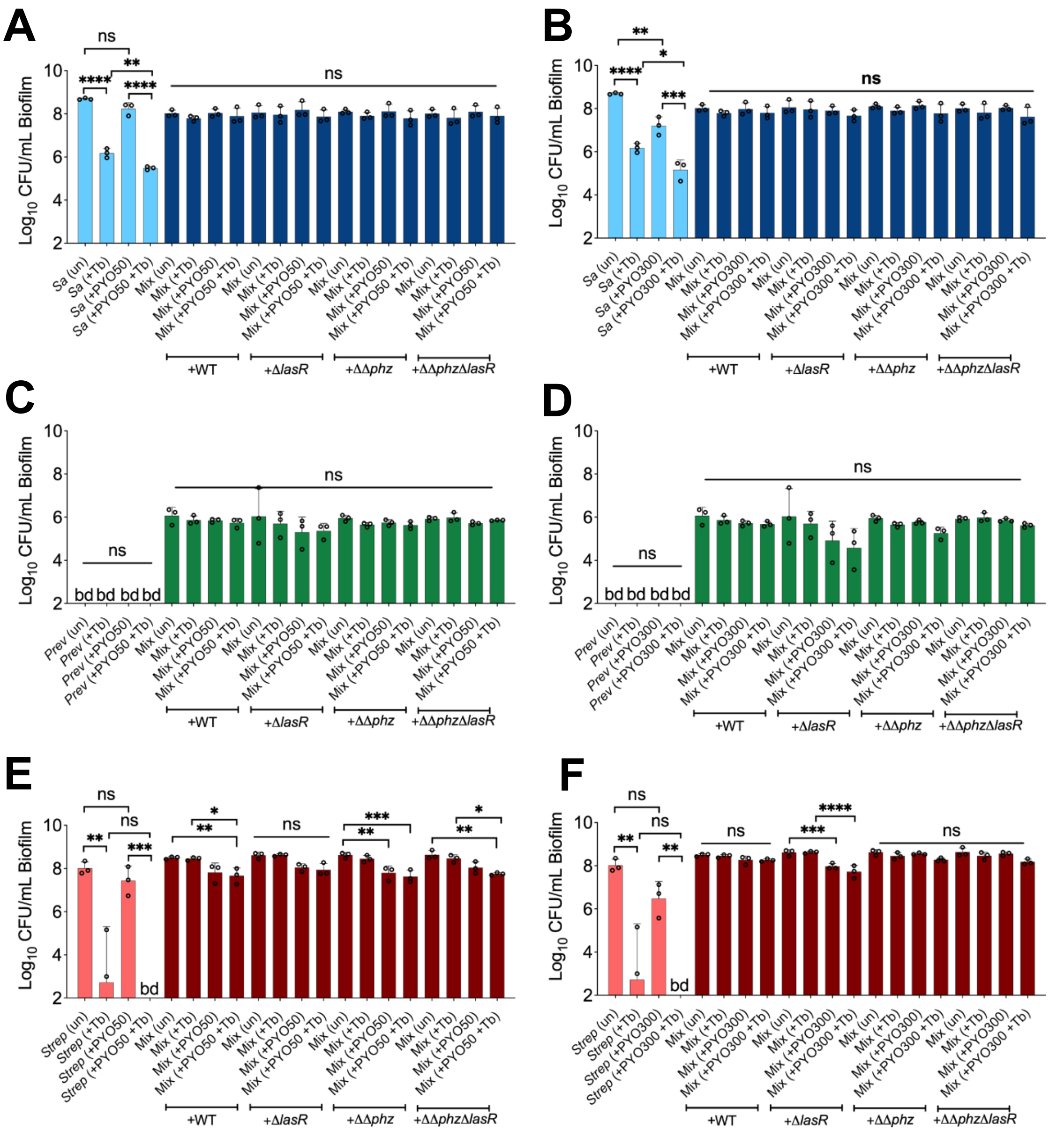

**Fig. S15. Impact of phenazine on community members treated with tobramycin: added phenazines and *P. aeruginosa* mutants do not impact other community members.** Colony forming units (CFUs) counts of *S. aureus* (*Sa*), *S. sanguinis* (*Strep*) and *P. melaninogenica* (*Prev*) grown as monoculture (light color) or mixed biofilm communities (dark color) co-cultivated with *P. aeruginosa* (WT) and associated mutants treated with tobramycin and (**A**, **C**, **E**) 50 µM (+PYO50) or (**B**, **D**, **F**) 300 µM (+PYO300) of phenazine. Each column represents the average from at least three biological replicates, each with at least three technical replicates. Statistical analysis was performed using ordinary one-way analysis of variance (ANOVA) and Tukey's multiple comparisons posttest with *, $P < 0.05$; **, $P < 0.01$; ***, $P < 0.001$, and ****, $P < 0.0001$, ns = non-significant, bd = below detection, un = untreated +Tb = +100 µg/mL tobramycin.

**Table S1. Minimal Bactericidal Concentration (MBC) of *P. aeruginosa* PA14 planktonic and biofilm cells treated with tobramycin exposed to the following conditions for 24 hrs.**

| **Condition** | **MBC** |
|---|---|
| ASM + tobramycin | 62.5 μg/mL |
| PA14 monoculture supernatant + tobramycin | 62.5 μg/mL |
| Δ*lasR* monoculture supernatant + tobramycin | 62.5 μg/mL |
| Community supernatant (WT) + tobramycin | 62.5 μg/mL |
| Community supernatant (Δ*lasR*) + tobramycin | 62.5 μg/mL |

**Table S2. Strains used in the study**.

| Species and strain | Strain number | Phenotype/Genotype | Ref. |
| --- | --- | --- | --- |
| ***P. aeruginosa*** | | | |
| PA14 | SMC232 | Laboratory reference strain | (10) |
| PA14Δ*lasR* | SMC5021 | in-frame deletion of *lasR* gene | (4) |
| PA14Δ*lasR*::*lasR* | SMC9421 | SMC5021 with complementation of *lasR* at the native locus | This study |
| PA14Δ*mvfR* | SMC5018 | In-frame deletion of *mvfR* gene | (11) |
| PA14Δ*rhlR* | DH2742 | In-frame deletion of *rhlR* gene | (12) |
| PA14ΔΔ*phz* | SMC5020 | In-frame deletions of *phzA1-G1* and *phzA2-G2* genes | (13) |
| PA14Δ*mvfR*Δ*lasR* | DH1111 | In-frame deletion of *lasR* and *mvfR* genes | (11) |
| PA14Δ*lasR*Δ*rhlR* | DH2944 | In-frame deletion of *lasR* and *rhlR* genes | (12) |
| PA14ΔΔ*phz*Δ*lasR* | SMC9422 | In-frame deletion of *phzA1-G1*, *phzA2-G2* and *lasR* genes | This study |
| NC-AMT0101-1-2 | DH2417 | Chronic lung infection isolate with functional LasR allele, parent of NC-AMT0101-1-1 | (14) |
| NC-AMT0101-1-1 | DH2415 | Chronic lung infection isolate related to DH2417 with LasR loss-of-function (frame shift) allele | (14) |
| PA14Δ*lasI*Δ*rhlI* | DH242 | In-frame deletions of *lasI* and *rhlI* genes | (15) |
| PAO-MW1qsc102 | DH161 | PAO1 Δ*lasI*Δ*rhlI* AHL-sensing *lacZ* bioreporter for 3-oxo-C12-HSL production | (16) |
| Clinical isolate | SMC1587 | Mucoid CF isolate | (17) |
| Clinical isolate | SMC1595 | Non-mucoid CF isolate | (17) |
| Clinical isolate | SMC1596 | Non-mucoid CF isolate | (17) |
| ***S. aureus*** | | | |
| Newman | SMC1007 | Methicillin susceptible *Staphylococcus aureus* | (18) |
| JE2 | SMC8668 | Methicillin resistant *Staphylococcus aureus* | (19) |
| USA300 | SMC6979 | Methicillin resistant *Staphylococcus aureus* | (20) |
| ***Streptococcus* spp.** | | | |
| *S. sanguinis* | SMC7474 | Strain SK36 | (21) |
| *S. constellatus* | SMC7155 | *Streptococcus milleri* group | (22) |
| *S. intermedius* | SMC7156 | *Streptococcus milleri* group | (22) |
| *S. anginosus* | SMC5342 | *Streptococcus milleri* group | (22) |
| ***Prevotella* spp.** | | | |
| *P. melaninogenica* | SMC6965 | ATCC25845 | (23) |
| *P. intermedia* | SMC5371 | ATCC25611 | (23) |

| | | | |
|---|---|---|---|
| ***E. coli*** | | | |
| SM10 λpir | SMC32 | Used as a conjugation partner for introducing pEX18-based plasmids. | |
| S17 λpir | SMC117 | Used as a conjugation partner for introducing pMQ30-based plasmids. | |
| **Plasmids** | | | |
| pEX18Gm-Δ*lasR* | DH123 | PA14 *lasR* in-frame deletion construct; $Gm^R$ | (4) |
| pMQ30 + *lasR* | DH3548 | For complementing WT *lasR* gene at the native locus; $Gm^R$ | (5) |
